# Allometric analysis reveals distinct anatomical patterning and genetic architecture across wheat nodal roots

**DOI:** 10.64898/2026.11.16.766962

**Authors:** Akshay B. Tawale, Matías Schierenbeck, Dylan H. Jones, Riccardo Fusi, Ivan Lopez-Valdivia, Annegret Wolf, Petra Linow, Daniele Sandoni, Aurélien D’Oria, Mustapha Arkoun, Juan I. Dietz, Pablo E. Campos, Andreas Börner, Guy Golan, Hannah M. Schneider

## Abstract

Root anatomical traits influence resource acquisition, metabolic costs, hydraulic conductance, and stress adaptation in cereal crops. Although classified as a single root class, nodal roots may differ anatomically according to their position along the culm. As shown in maize, this variation is particularly notable in root cross-sectional area (RxA). However, it remains unclear whether these differences reflect developmental differentiation or simply variation associated with RxA. We quantified root anatomical traits across four main-culm nodes (basal to apical) in 190 hexaploid wheat genotypes grown under field and greenhouse conditions. To distinguish RxA-dependent from RxA-independent anatomical variation, we applied allometric size correction, adjusting traits to a reference RxA, and conducted parallel genome-wide association studies (GWAS) using raw and size-corrected traits. Raw measurements revealed increases in RxA and cortical area from basal to apical nodes, accompanied by decreases in metaxylem area. After correction to a common reference RxA within each environment, anatomical differences among nodal positions persisted, with roots emerging from more apical nodes exhibiting smaller stele and metaxylem areas. Differences in cortical area were primarily associated with greater cortical cell file layer number rather than increased mean cortical cell diameter. GWAS identified largely distinct genetic associations for raw and RxA-corrected anatomical traits, varying across nodal positions and environments. Together, these findings demonstrate that anatomical variation across the wheat nodal root system cannot be explained by RxA alone. Our results highlight nodal position and allometric scaling as important considerations for anatomical phenotyping and distinguishing genetic associations with overall root cross-sectional area from those with tissue organisation.

**Graphical abstract:** 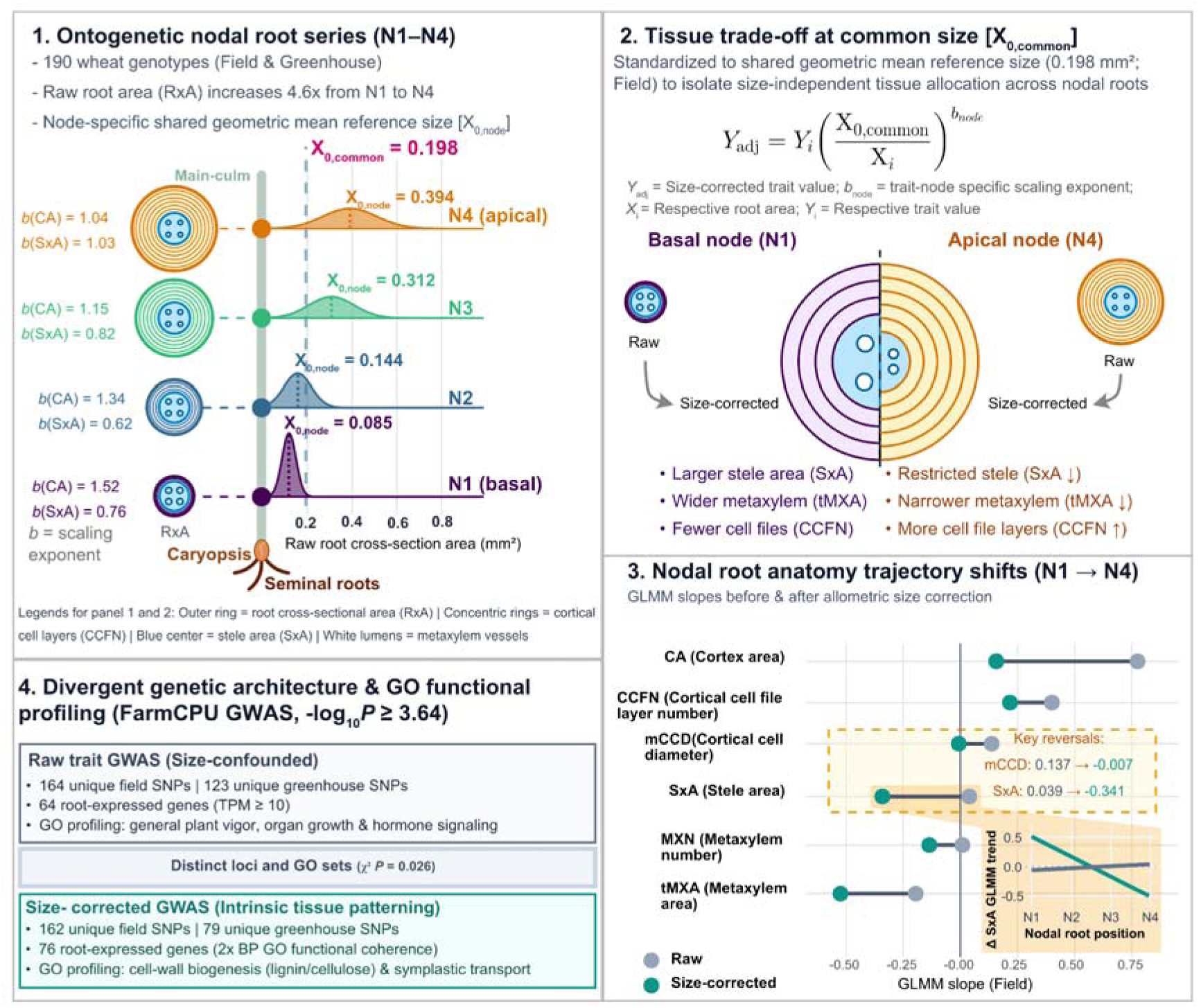

## 1. Introduction

Wheat (*Triticum aestivum*) is one of the most important cereal crops, providing approximately 20% of global dietary calories (*1*). It is the third most widely produced cereal after maize and rice, and remains a major component of human diets worldwide. Maintaining wheat productivity under changing climates and increasing resource limitations will require a better understanding of the traits that determine resource acquisition and stress adaptation (*2*). Roots are central to these processes because they acquire water and nutrients, anchor the plant to the soil, and mediate responses to edaphic stress (*3*). Despite these functions, root traits remain comparatively underexplored, in part because techniques for phenotyping belowground organs are relatively underdeveloped compared to aboveground tissues, and limited sample accessibility in field conditions makes root studies especially challenging (*4*). Wheat, like most grasses, develops a fibrous root system composed of roots with distinct developmental origins: embryonic (also called seminal) roots that develop from the embryo and post-embryonic (or nodal) roots that emerge from culm nodes. Roots emerging together from a single culm node are referred to collectively as nodal root cohorts. Both root types subsequently produce root-borne lateral roots (*5*, *3*). In wheat and other cereals, post-embryonic nodal roots develop from basal nodes of the main culm and tillers (*6–8*). These basal nodes remain tightly compressed due to minimal internode elongation, with successive phytomer lengths increasing non-uniformly along the main culm axis (*9*). However, despite originating in close physical proximity along the culm, successive nodal root cohorts appear sequentially throughout plant development, driven by thermal time accumulation (*10*).

While the embryonic root system dominates early development in wheat and barley (*Hordeum vulgare*) (*11–13*), nodal roots can make an increasingly large contribution to water and nitrogen acquisition as plant matures (*14–18*). For example, a hydroponic study of wheat, barley, and rye (*Secale cereale*) found that between anthesis and physiological maturity, nodal roots accounted for approximately 75% of root water uptake, with seminal roots contributing the remaining 25% (*19*). Under hypoxic or waterlogged conditions, wheat nodal roots can maintain greater nutrient uptake and remain functional longer than seminal roots (*20*). Together, these findings highlight the increasing importance of nodal roots as plants mature and their critical role in sustaining resource acquisition under both favourable and stressful conditions (*15*, *17*, *18*, *21*).

While the contribution of nodal roots to resource acquisition changes as plants mature, their functional capacity is also determined by their internal anatomical organisation. Root anatomical traits influence the metabolic cost of soil exploration, mechanical properties, and the transport of water and nutrients (*22*, *23*). Plants can reduce the metabolic cost of root construction and maintenance through anatomical phenotypes, including reduced cortical area, increased cortical cell size, fewer cortical cell layers, and greater aerenchyma formation in the cortex. These traits can reduce respiratory costs and enable greater soil exploration (*22*). Increased root diameter can prevent buckling, facilitate penetration in hard soils, and support plant anchorage (*24–26*). Root anatomy also regulates hydraulic function: fewer and narrower metaxylem vessels can reduce axial hydraulic conductance, potentially limiting early-season water use and conserving soil water for later reproductive development, and tissues in the cortex, including aerenchyma, can influence radial transport of water and nutrients (*27*, *28*). Thus, variation in root anatomy can have direct consequences for both resource acquisition and whole-plant performance (*29–31*).

Importantly, root anatomy is not uniform across the cereal root system. Anatomical traits differ between embryonic and nodal root classes in cereals such as maize, barley, and intermediate wheatgrass (*Thinopyrum intermedium*) (*21*, *32*, *33*). These root classes have distinct developmental origins and can differ in their genetic regulation (*7*). However, anatomical variation may not be limited to differences between embryonic and post-embryonic roots. Evidence from maize indicates that nodal roots originating from successive culm nodal positions differ systematically in their anatomy and architecture; in particular, apical nodal roots typically exhibit larger cross-sectional, cortical, and stelar areas than roots from basal nodes (*33–35*).

Such variation is particularly relevant because cereals produce nodal roots sequentially from multiple culm nodes as the plant matures; typically, up to nine nodes in maize and three to seven in wheat (*36*, *37*, *10*). Root cohorts emerging from successive culm nodes therefore represent a structured developmental series rather than repeated, functionally equivalent units of a single root class. Similar positional differentiation occurs aboveground, where leaves and internodes associated with successive phytomers differ systematically in their morphology and function, for example, the flag leaf in wheat (*36*, *38*). Thus, culm node position may provide a developmental coordinate associated with anatomical and potentially functional differentiation within the cereal root system. Consequently, treating all nodal roots as a single root class, or assuming that anatomical measurements of early-emerging basal roots represent those produced later in plant development, may obscure developmental variation, particularly when environmental conditions change over time (*39*).

A major challenge, however, is determining whether anatomical differences in root cohorts across nodal positions reflect developmental differentiation or simply differences in root cross-sectional area. Root tissues commonly exhibit allometric relationships, in which the dimensions of individual tissues change predictably as total root cross-sectional area increases (*34*) Thus, without isolating this RxA-dependent scaling, it is not possible to determine whether successive nodal cohorts follow distinct developmental blueprints for tissue organisation or are simply larger versions of the same developmental blueprint. Distinguishing RxA-dependent scaling from RxA-independent developmental differentiation is therefore essential to accurately phenotype variation across the nodal root system.

Resolving this dependence of anatomical traits on RxA is equally important for uncovering the true genetic architecture of root anatomy. Without correction, genetic associations mapped to anatomical traits may partly reflect variation in RxA rather than tissue organisation alone. Furthermore, conventional normalisation approaches, such as simple ratios or linear residual correction, may not adequately account for nonlinear scaling relationships between RxA and individual tissues, potentially introducing statistical artefacts or obscuring biological associations (*40*). Size correction provides an alternative by estimating trait-specific scaling relationships and adjusting anatomical measurements to a common RxA (*41–44*). By applying this framework across genotypes and nodal positions, we can distinguish genetic associations with overall root cross-sectional area from those associated with tissue organisation after accounting for RxA (*44–46*). However, it remains unclear how wheat root anatomy varies across successive nodal positions, how much of this variation is explained by allometric scaling, and whether raw and size-corrected anatomical traits yield distinct genetic architectures.

We hypothesised that anatomical variation across successive nodal positions in wheat reflects developmental differentiation, extending beyond mere differences in root cross-sectional area. To test this hypothesis, we characterised root anatomy across four nodal cohorts in a diversity panel of 190 hexaploid wheat genotypes grown under field and greenhouse conditions. Because total root cross-sectional area (RxA) represents the primary physical boundary constraining tissue expansion, we quantified raw anatomical traits and their allometric relationships with RxA, subsequently applying allometric size correction to distinguish RxA-dependent from RxA-independent variation. Throughout this study, “size-corrected” traits refer to anatomical measurements adjusted to a common reference RxA. We then conducted parallel genome-wide association studies (GWAS) on both raw and size-corrected anatomical traits to identify genetic loci associated with overall anatomical size versus intrinsic tissue organisation. Together, these analyses provide a framework for characterising root anatomical differentiation across nodal positions while explicitly accounting for allometry. Because culm nodal position is inherently associated with differences in root emergence time, root age, and whole-plant developmental stage, we interpret this anatomical variation within a broader developmental context rather than as a rigid, isolated effect of the node itself.

## 2. Materials and methods

### 2.1. Plant material

A genetic diversity panel of 190 hexaploid winter and spring wheat (*Triticum aestivum* L., 2n = 6x = 42) genotypes, designated the Tango-Wheat Panel, was used in this study. This panel was structured to represent modern elite germplasm, primarily adapted to temperate agricultural zones. It comprises 157 commercial cultivars released in Argentina between 2000 and 2024, one from Brazil, 25 from Uruguay, two from Germany, and five elite lines from the International Maize and Wheat Improvement Centre (CIMMYT, Mexico). Detailed information, growth habit (days from emergence to flag leaf expansion (GS39) and days from emergence to anthesis initiation (GS61)), and origin for the entire panel are compiled in Table S1.

### 2.2. Field trial

The Tango-Wheat panel was grown during the spring season (March 2025 to July 2025) on the experimental field of the Leibniz Institute of Plant Genetics and Crop Plant Research (IPK), Gatersleben, Germany (51°49’22.0” N 11°16’13.2” E). The experimental soil type is Chernozem, characterised by a sandy clay loam texture (*47*). Seeds were sown using an automated experimental seeder at a standardised density of 300 seeds m^−2^. The trial was laid out as a flat-block design with 190 individual genotypes, each allocated to a single plot measuring 1.2 m² (1.2 m length × 1.0 m width) with six longitudinal rows spaced 0.2 m apart. Perimeter buffer plots containing the commercial cultivar ‘KWS Quintus’ were sown around the entire trial boundary to minimize edge effects.

The crop was managed using conventional tillage and standard agronomic practices. Nitrogen fertiliser was applied as urea in three split applications of 60 kg N ha^-^¹ each: at the onset of the vegetative period (GS13, (*48*)), during stem elongation (GS31/32), and at the booting stage (GS47–49), for a total seasonal nitrogen application of 180 kg N ha^-^¹. Chemical weed control was achieved using 5 g a.i. ha^-^¹ florasulam + 45 g a.i. ha^-^¹ pinoxaden (Axial komplett®) and 20 g a.i. ha^-^¹ carfentrazone-ethyl during post-emergence for grass weed control, and eight days after using 700 g a.i. ha^-^¹ mecoprop-P + 320 g a.i. ha^-^¹ 2,4-D (Aurora 40 WG® and Duplosan KV neu®) for broadleaf weed control. Deltamethrin was applied at 7.5 g a.i. ha^-^¹ (Decis Forte®) to control insect pests. The entire field trial was grown under strictly rainfed conditions. Daily meteorological data, including ambient temperature, precipitation, and relative humidity, were recorded by a dedicated weather station located 200 m from the plots (Figure S1).

### 2.3. Greenhouse experiment

A parallel experiment was conducted in the greenhouse facilities of IPK Gatersleben. Sowing was performed 3 cm deep in individual cells of 24-well Quickpot trays (model 24T/16), with one seed per cell, following a randomised complete block design (RCBD) across three independent blocks (replications). Each growth cell measured 55 × 60 × 160 mm (volume ∼330 cm³). The growth substrate consisted of sand (particle size 0.20–0.63 mm), pre-mixed with Osmocote® Pro 3-4M controlled-release fertiliser (19% N, 9% PCOC, 10% KCO + 2% MgO + micronutrients; 3 to 4-month release profile) at a concentration of 8 g L^-^¹ of sand.

Plants were maintained under a 16h light / 8h dark photoperiod. During the light period, photosynthetic photon flux density of 900 µmol m^-^² s^-^¹ was provided at the canopy level. Air temperature was regulated at 17 °C during the day and 15 °C at night. Root samples were collected 8 weeks after sowing (960 GDD), once plants had reached the booting stage.

### 2.4. Root sampling and anatomical phenotyping

Root systems in both greenhouse and field environments were targeted for harvest at the booting stage (GS45–49). To account for genotype-specific phenological variation, field sampling dates were adjusted within a ±2-week window. Following the shovelomics approach (*49*, *50*), field root crowns were excavated from rows 2 and 5 of each 6-row plot to ensure uniform stand density and avoid edge effects (*51*). Excavated root crowns were soaked in a dilute liquid-soap solution for 18 h to loosen soil aggregates, then gently rinsed with a low-pressure water spray. Three representative plants were selected as biological replicates (Figure 1). In parallel, greenhouse-grown plants were extracted from Quickpot trays and washed sand off the root system with a gentle water spray.

**Figure 1.**
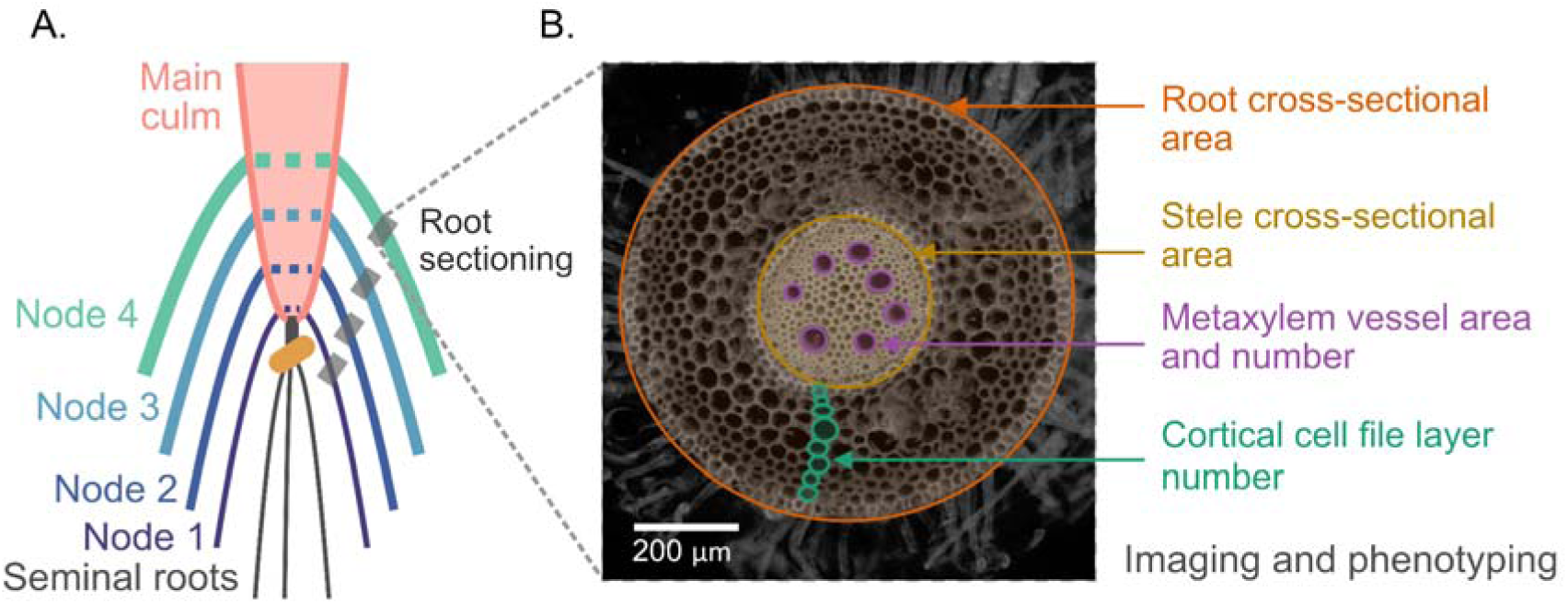
Nodal root sampling strategy and anatomical phenotyping. (A) Schematic of the main culm illustrating distinct nodal root classes (Nodes 1–4) and the sectioning zone, at 2 cm distal to the root-shoot junction. (B) Representative transverse root section highlighting the quantified anatomical traits: root cross-sectional area, stele cross-sectional area, metaxylem vessel area and number, and cortical cell file layer number.

The main culm root system was preserved in 70% ethanol at 4 °C until further sub-sampling for the anatomical analysis. Main-culm nodal roots originating from the first four culm nodes (N1 to N4, where N1 = oldest/most basal node, node 1, and N4 = youngest/most apical node, node 4) were sampled sequentially. Transverse sections were cut precisely at a distance of 2Ccm from the root-shoot emergence junction (*39*, *52*, *53*). The theoretical maximum sampling size comprised 4,560 anatomical specimens (2 growth conditions × 190 genotypes × 4 nodal positions × 3 biological replicates). We obtained 4242 anatomical specimens in total for further image analysis (Table S2).

Sampled roots were mounted on the Rapid Anatomics Tool and imaged without chemical staining using UV-induced cell-wall autofluorescence (excitation: 340-380 nm; emission: >420 nm). Digital RGB images were acquired in PNG format using a digital microscope camera and Dino-Capture 2.0 (v1.553B) software (*54*). Magnification was set to 190x-220x for basal nodes (N1 and N2) and 175x-190x for apical nodes (N3 and N4) to account for increasing root cross-sectional area (Figure 1).

Image processing and trait quantification were performed in Fiji (ImageJ v2.16.0) (*55*). Captured RGB images were converted to 8-bit, and ten spatial, meristic, and relational anatomical traits were measured manually using region-of-interest (ROI) tools (Figure 1, Table 1).

**Table 1.**
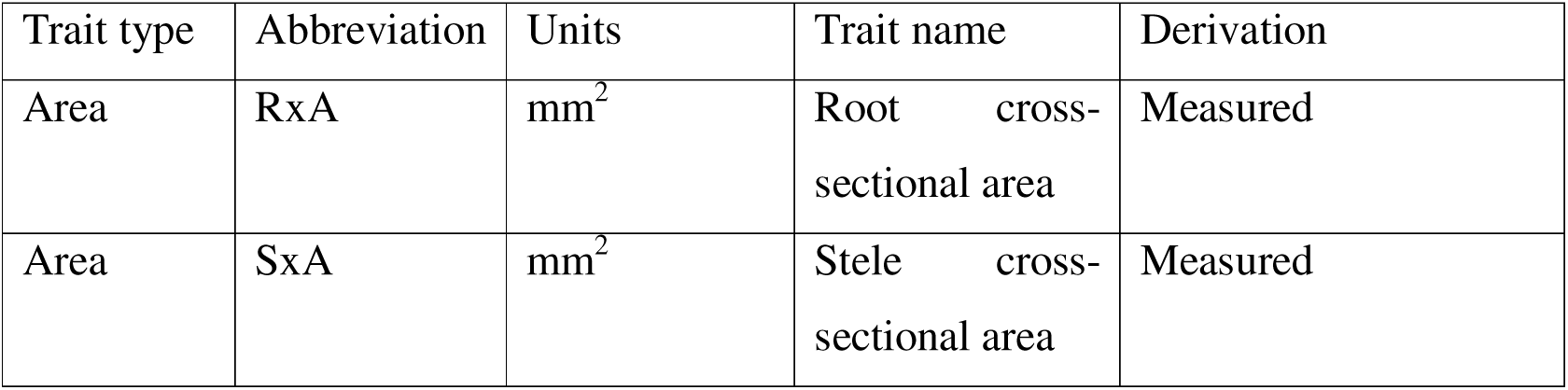
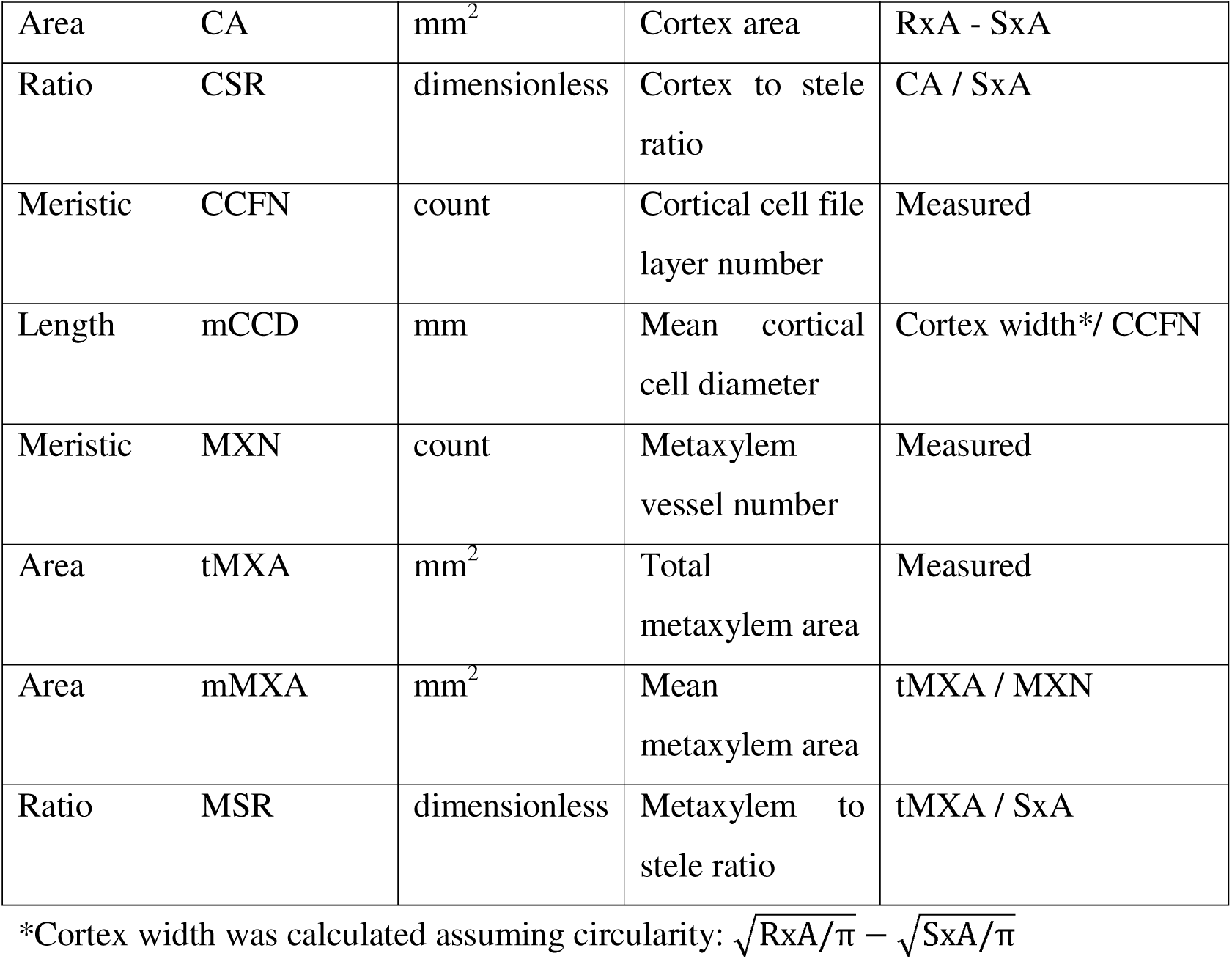
Description of wheat root anatomical traits measured across nodal positions.

| Trait type | Abbreviation | Units | Trait name | Derivation |
|---|---|---|---|---|
| Area | RxA | mm<sup>2</sup> | Root cross-sectional area | Measured |
| Area | SxA | mm<sup>2</sup> | Stele cross-sectional area | Measured |
| Area | CA | mm<sup>2</sup> | Cortex area | RxA - SxA |
| Ratio | CSR | dimensionless | Cortex to stele ratio | CA / SxA |
| Meristic | CCFN | count | Cortical cell file layer number | Measured |
| Length | mCCD | mm | Mean cortical cell diameter | Cortex width*/ CCFN |
| Meristic | MXN | count | Metaxylem vessel number | Measured |
| Area | tMXA | mm<sup>2</sup> | Total metaxylem area | Measured |
| Area | mMXA | mm<sup>2</sup> | Mean metaxylem area | tMXA / MXN |
| Ratio | MSR | dimensionless | Metaxylem to stele ratio | tMXA / SxA |

\*Cortex width was calculated assuming circularity: $\sqrt{\mathrm{RxA}/\pi} - \sqrt{\mathrm{SxA}/\pi}$

### 2.5. Allometric modelling and size-correction

#### 2.5.1. Bivariate allometric modelling

To distinguish RxA-dependent from RxA-independent anatomical variation, bivariate scaling relationships between individual traits (*Y*) and root cross-sectional area (RxA, *x*) were modelled using Huxley’s power equation (*56*):

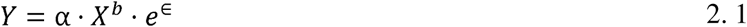

α is the allometric coefficient, *b* is the scaling exponent, and e^□^ represents multiplicative error. For statistical estimation of scaling exponents, variables were linearised into base-10 logarithmic space:

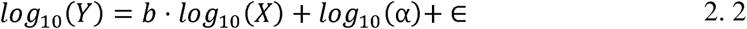

We used a hybrid analytical protocol to address distinct biological and mathematical objectives (*44*). Standardised Major Axis (SMA) regression was used strictly to characterise physiological scaling relationships because it accounts for biological variance in both variables. However, because SMA slopes inflate when trait-size correlations are weak (*b*_SMA_ =*b*_OLS_/∣r∣), applying them directly to size-correction formulas yields unstable, over-corrected phenotypes for low-R^2^ traits (e.g., metaxylem-related traits, Table S4) (*57*).

Therefore, the mathematical size-correction step utilised trait-specific statistical models at the replicate level to accurately estimate the correction exponent (*b*). Continuous spatial traits were modelled using ordinary least squares (OLS) regression in log_10_-log_10_ space. Strictly positive count variables (MXN, CCFN) were treated as discrete count traits rather than categorical variables and modelled using Poisson or negative binomial generalised linear models (GLMs) fitted directly to raw counts, with the resulting slope used as the allometric scaling exponent. Although ratio traits (CSR, MSR) are already expressed relative to another anatomical measurement, they may still vary with RxA. Therefore, ratio traits were not scaled directly but were computed post-hoc from their independently size-corrected spatial components.

#### 2.5.2. Slope homogeneity

Before standardisation, an analysis of covariance (ANCOVA) was executed for each environment-trait to formally test for slope homogeneity across nodal positions using the model:

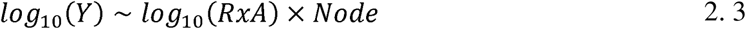

A significant interaction term (*log_10_(RxA) x Node*, *P* < 0.05) confirmed heterogeneity in scaling slopes across nodes (e.g., in a field dataset, CA scaling shifts from hyperallometry [b=1.52] at Node 1 to isometry [b=1.04] at Node 4). Although some slope values were more uniform in the greenhouse dataset, we utilised node-specific slope values to maintain analytical consistency across both datasets (Figure 3, Table S3).

#### 2.5.3. Allometric size correction

##### 2.5.3.1. Node-specific allometric size correction for GWAS

To preserve the biological identity and ontogenetic context of each distinct nodal root position, we used an environment- and node-specific common within-group standardisation strategy. This node-specific reference baseline area was estimated using a geometric mean of RxA (x_0,node_ (mm^2^); N1 = 0.085, N2 = 0.144, N3 = 0.312, N4 = 0.394; for the field dataset; Fig. S6) for roots from that specific nodal position and environment. Trait values were then standardised (or size-corrected) to a node-specific baseline area using the allometric scaling equation (*44*) adapted to our case scenario:

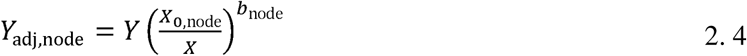

Transformed to a linear logarithmic format, the equation is expressed as:

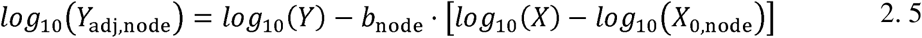

where *Y*_adj,node_ is the size-corrected phenotype, *Y* is the raw trait value, *x* is the raw individual RxA at which the *Y* trait value is measured, and *x*_0,node_ is the reference geometric mean RxA for a given node. The scaling exponent, *b*_node_, was fitted independently for each combination of trait, node, and environment.

##### 2.5.3.2. Common referenced size correction for physiological/developmental comparisons

To facilitate the physiological comparison of anatomical trait trajectories (i.e., the sequential patterns of change in a given trait across successive nodal cohorts from N1 to N4), a common reference root cross-sectional area (RxA) within each environment is required. To avoid the mathematical distortions of scaling roots outside their natural biological ranges, we identified the range of overlapping RxA values among nodal positions separately for each environment (0.09 to 0.39 mm^2^; field) and set the common reference RxA (*x*0,common (mm^2^); 0.198 for field and 0.699 for greenhouse dataset; Figure S7) to the grand geometric mean RxA. Traits were standardised using the node-specific scaling exponents (*b*_node_) projected to this shared baseline within each environment.

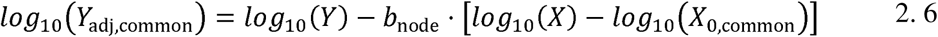

This allometric adjustment reduces the dependence of anatomical traits on RxA, and the resulting traits are referred to as size-corrected traits for physiological comparisons. Thus, node-specific reference RxA values were used for within-node genetic analyses, whereas a common reference RxA within each environment was used to compare anatomical traits across nodal positions.

##### 2.5.3.3. Empirical validation and quality diagnostics for the size correction protocol

To validate whether the standardisation pipeline has effectively removed the dependence of stele and cortical traits on RxA in nodal roots, we did two trait-specific diagnostic checks. Post-correction residual correlations (*r*) between size-corrected traits and RxA were reduced to near zero (|*r*| ≤ 0.24) and were non-significant (*P* ≥ 0.05) for the vast majority of combinations, confirming the removal of size-dependent confounding effect. Furthermore, Spearman rank correlations (ρ) between raw and size-corrected entry means demonstrated rank preservation (ρ ≥ 0.75, *P <* 0.001) for metaxylem-related traits (mMXA, MSR, MXN, tMXA) and mean cortical cell diameter (mCCD). In contrast, cortical area (CA) demonstrated near-complete rank shuffling across all nodes (ρ ranging from -0.30 to 0), mathematically confirming that phenotypic variation in unadjusted CA is almost entirely driven by whole-root scaling rather than independent tissue partitioning (Figure S8).

### 2.6. Generalised Linear Mixed-Effects Models (GLMMs) for trait trends

Raw trait trajectories across nodes N1 to N4 were modelled using glmmTMB (*58*) fitted on replicate-level data:

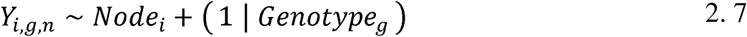

where *Y*_i,g,n_ is the phenotypic value of replicate *n* of genotype *g* at nodal position *i (Node*_i_; fixed effect, N1–N4), and (1 | *Genatype_g_*) represents the random intercept for genotype. Error distributions were assigned by trait family: Gamma (log link) for the area and ratio traits (RxA, CA, CSR, SxA, tMXA, mMXA, mCCD); Poisson (log link, auto-upgrading to Negative Binomial if overdispersed) for count traits (MXN, CCFN); and Beta (logit link) for bounded proportion traits (MSR). Pairwise node differences were evaluated via Tukey’s HSD test (*P* < 0.05) within each trait.

### 2.7. Genotypic BLUE calculations, genotypic repeatability (*R*) and GWAS

#### 2.7.1. Environment-specific BLUE methodology

Genotypic Best Linear Unbiased Estimators (BLUEs) were calculated independently for every environment-node-trait combination. For the field, as single-plot layouts confound plot and genotype, BLUEs were calculated as the arithmetic mean of the three within-plot plant subsamples per genotype:

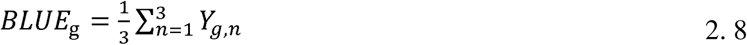

For the randomised complete block design (RCBD) in the greenhouse experiment, block-level variance was removed by fitting a fixed-effects linear model per node-trait combination:

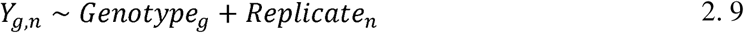

Where *Genatype_g_* is the fixed genotypic effect, and *Rep1icate*_n_ is the fixed block effect. Adjusted marginal means were extracted as BLUEs using emmeans (v2.0.4).

#### 2.7.2. Genotypic repeatability (*R*)

Environment-specific genotypic repeatability (*R*) was estimated independently for each trait-node combination using both raw unadjusted (*Y*) and node-specific, size-corrected (*Y*_adj,node_) trait values. Variance components were extracted using random-effects models fitted with the lme4 package (*59*), matching the model structures used for the BLUE estimations described above. Repeatability was calculated on an entry-mean basis, accounting for unbalanced data (*60*) as:

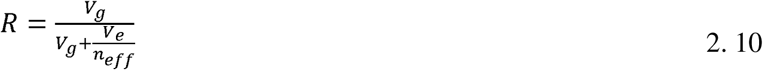

Where *v_g_* and *v_e_* are the genotypic and residual variances, respectively, and *n_eff_* is the harmonic mean of replicates per genotype to adjust for unbalanced replication. Models that produced a singular fit (where a variance component is estimated at the zero boundary) were flagged but retained, as the resulting *R estimates remain mathematically valid for quantitative scaling (Figure S13)*.

We conducted two parallel GWAS analyses per trait, node (N1–N4) and environment combination. GWAS using size-corrected traits was used to identify genetic associations with anatomical variation after accounting for RxA, whereas GWAS using raw traits identified associations with unadjusted anatomical variation, including variation associated with RxA.

#### 2.7.3. Genotyping and Genome-Wide Association Studies (GWAS)

The Tango-Wheat panel was genotyped by TraitGenetics (SGS Institut Fresenius GmbH, Gatersleben, Germany). For genomic DNA extraction, leaf tissue samples were taken from young seedlings for each accession. The genotyping was performed using the 25K Infinium iSelect array, providing 24,243 SNP markers after quality filtering (minor allele frequency MAF > 0.05, missingness < 10%).

Genome-wide association studies were conducted in R using the Genome Association and Prediction Integrated Tool (GAPIT v3.0; (*61*)) with both the Multiple Loci Mixed Model (MLMM; (*62*)) and the Fixed and Random Model Circulating Probability Unification (FarmCPU; (*63*)) models. MLMM was used for model validation.

To assess potential confounding factors, population structure and pairwise kinship were characterised via SNP marker PCA (first four PCs) and the VanRaden matrix (VanRaden, 2008), respectively. To define marker-trait associations across all GWAS runs, a genome-wide threshold was established using LD-based marker pruning in PLINK v1.9 (r^2^ = 0.2). This retained M_e_ = 4,402 independent SNPs out of 24,243 total markers, establishing a suggestive genome-wide threshold of 1/M_e_ (*P* = 2.27E-4, i.e., −log_10_ *P* ≥ 3.64) (*64*).

### 2.8. Gene Ontology (GO) enrichment analysis and multi-omics candidate gene prioritisation

To identify overrepresented genomic pathways among genes that met the suggestive threshold, we performed Gene Ontology (GO) enrichment analysis. The gene stable IDs and GO terms were retrieved from the Ensembl Plants BioMart database. To control for genomic coverage biases inherent in SNP arrays and prevent false-positive enrichment, a local background universe was established comprising all high-confidence wheat genes (IWGSC RefSeq v2.1) located within a ±200 kb window of any tested SNP (*65*). GO enrichment was performed using a hypergeometric test via the enricher function in the R package clusterProfiler (*66*). Enriched terms were classified into “Plant vigour/growth” or “Tissue allocation/patterning” via keyword matching (Table S11), and functional divergence between the size-corrected and unadjusted raw models was evaluated using Chi-square tests.

To further identify candidate genes associated with loci detected in the GWAS, we evaluated SNPs that exceeded the suggestive threshold (−log_10_ *P* ≥ 3.64) in the FarmCPU model. We used an LD-aware windowing approach, in which genome-wide LD decay was estimated to be ∼5.3 Mb (r^2^ = 0.2). Loci were initially defined using a core region of ±200 kb and an extended window of ±2 Mb surrounding each lead SNP to retrieve primary encoded proteins (*65*). Candidate genes were subsequently prioritized through orthology mapping in *Arabidopsis thaliana* and rice (*Oryza sativa*) using Ensembl Plants and Monocots PLAZA 5.0 (*67*, *68*), functional annotations from UniProt (UniProt Consortium; https://www.uniprot.org/), expression profiles from the Wheat eFP Browser (*69*, *70*), and documented functional associations with the target traits (*71*).

To specifically investigate root-specific expression patterns of candidate genes, we utilised the metadata table to annotate each run_accession with high-level plant organ designation: “roots”, “leaves/shoot”, “spike”, or “grain”. Expression data were downloaded. For each gene and organ category, the expression signal was calculated as the average TPM across the five samples with the highest expression within that organ. Furthermore, genes with a maximum organ-average TPM < 10 were excluded to retain only moderately to highly root-expressed candidates (requiring TPM ≥ 10) (*72*).

### 2.9. Statistical analysis

All data processing, statistical modelling, and visualisation were performed in R (v.4.6.0) using RStudio (*73*). Data wrangling and ingestion were done with the tidyverse suite (v2.0.0), while multivariate analysis [principal component analysis (PCA)] was done using FactoMineR (v2.18) and factoextra (v2.2.0). Mixed-effects models were fitted using lme4 (*59*) and glmmTMB (*58*); with residual diagnostics performed using DHARMa (v0.5.0); allometric scaling slopes were calculated using smatr (*74*) (continuous traits) and MASS (v7.3-66) (count traits, via Poisson/negative binomial regression); and genotypic BLUEs were calculated using emmeans (v2.0.4). Genome-wide association analyses were performed using GAPIT v3.0 (*61*). Finally, data visualisations and post hoc mean comparisons (Tukey’s HSD) were generated using ggplot2 (v4.0.3), along with extension packages such as patchwork (v1.3.2), ComplexUpset (v1.3.3), eulerr (v8.3.1), and multcompView (v0.1-12).

For downstream genomics, coordinate parsing and background universe construction were handled via GenomicRanges (v1.64.0) and rtracklayer (v1.72.0), while GO enrichment and comparative clustering were executed using clusterProfiler (*66*).

## 3. Results

### 3.1. Unadjusted root anatomy varies systematically across successive nodal positions

Before allometric correction for root cross-sectional area (RxA), root anatomical traits varied systematically among roots emerging from the first four nodes of the main culm (N1–N4; Figure 2). In the field, root cross-sectional area (RxA) increased from basal to apical nodal positions (GLMM slope = 0.531, *P <* 0.001). Cortical area (CA), cortical cell file layer number (CCFN), mean cortical cell diameter (mCCD), and cortex-to-stele area ratio (CSR) also increased (slopes = 0.775, 0.400, 0.137, and 0.707, respectively; all *P <* 0.001). Stele area (SxA) increased more modestly (slope = 0.039, *P <* 0.001). Metaxylem traits showed the opposite trend across nodal root cohorts from N1 to N4. Total metaxylem area (tMXA), mean metaxylem area (mMXA), and metaxylem-to-stele ratio (MSR) declined from N1 to N4 (slopes = −0.195, −0.210, and −0.261, respectively; all *P <* 0.001), whereas metaxylem vessel number (MXN) did not change significantly (slope = 0.00864). Consequently, increasing RxA across nodal positions was accompanied by greater cortical allocation and a modest increase in stele area, whereas metaxylem area declined (Figure 2).

**Figure 2.**
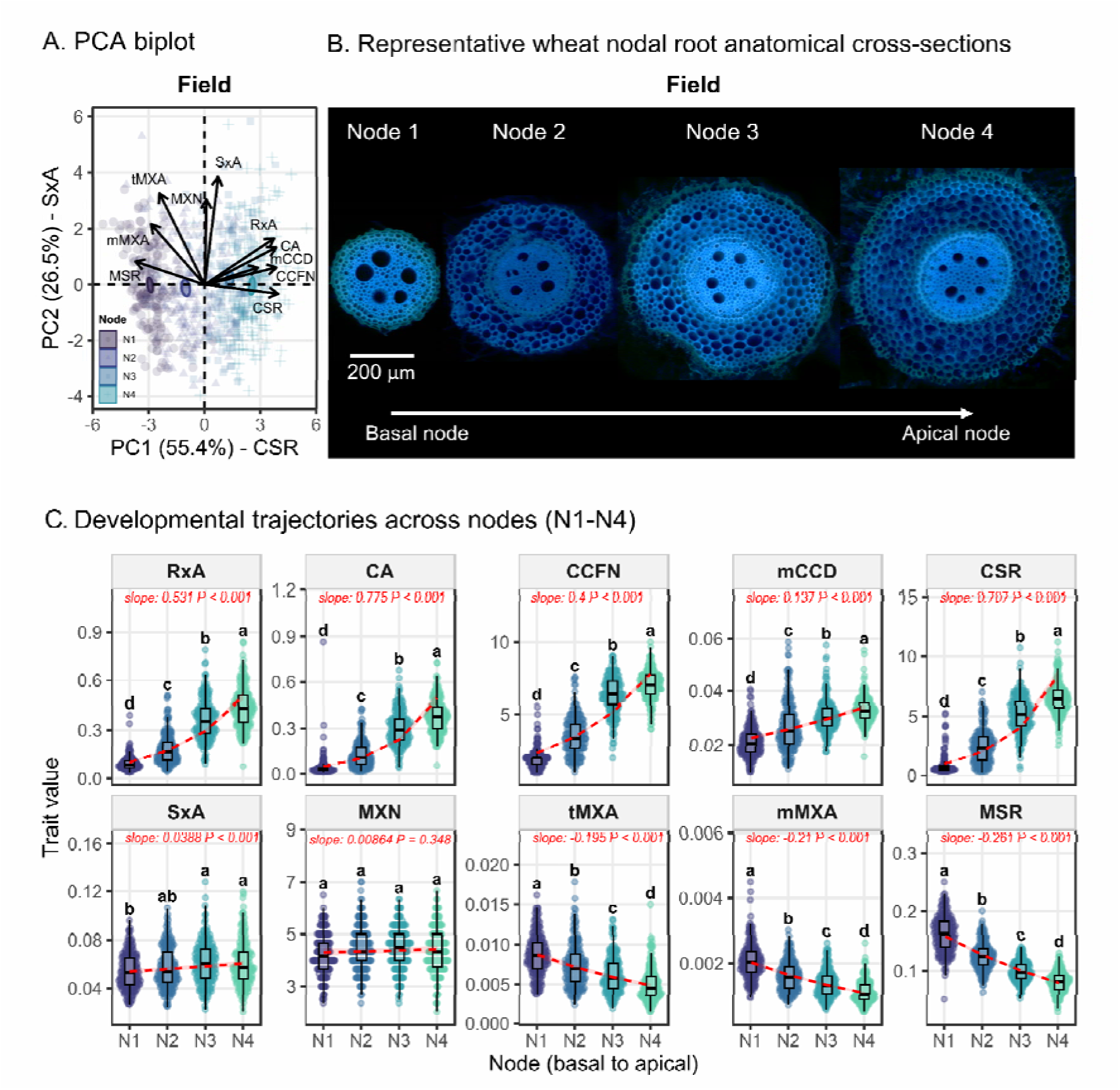
Anatomical variation across successive nodal positions in field-grown wheat. Root anatomical traits vary systematically among roots emerging from four successive nodes (N1–N4) along the main culm, from basal to more apical positions. (A) Principal component analysis (PCA) of anatomical traits across four nodal positions. Genotype means were centred and estimated before analysis. Points are coloured by nodal position, and ellipses represent 95% confidence regions. Arrows indicate trait loading vectors, and axes show the percentage of variance explained by each principal component. (B) Representative nodal root anatomical cross-sections at four nodal positions (Node 1 - 4; basal to apical nodes). (C) Variation in root anatomical traits across nodal positions, including root cross-sectional area (RxA), cortex area (CA), cortical cell file layer number (CCFN), mean cortical cell diameter (mCCD), cortex-to-stele area ratio (CSR = CA/SxA), stele cross-sectional area (SxA), metaxylem vessel number (MXN), total metaxylem area (tMXA), mean metaxylem area (mMXA), and metaxylem-to-stele area ratio (MSR = tMXA/SxA). Boxplots show the distribution of genotype means at each nodal position. Dashed red lines and shaded bands indicate predicted trends and 95% confidence intervals from generalised linear mixed-effects models (GLMMs) fitted to replicate-level data, with nodal position as a fixed effect and genotype as a random intercept. Gamma models with a log link were used for continuous traits, Poisson models with a log link for count traits, and a beta model with a logit link for MSR. Model slopes and associated P-values are shown in red. Different lowercase letters indicate significant differences among nodal positions (Tukey’s HSD, *P* < 0.05). The data represent 184 to 190 wheat genotypes, with each point corresponding to the mean of three biological replicates. A detailed summary can be found in Table S5.

### 3.2. Root cross-sectional area is strongly associated with cortical traits

Across individual nodal positions, RxA was strongly correlated with CA (Pearson r = 0.95-1.00) and moderately to strongly correlated with SxA (r = 0.48 - 0.86; Figure S3). Relationships between RxA and other cortical traits (CCFN, mCCD, and CSR) were more variable, while metaxylem traits generally showed weaker associations (r = −0.32 to 0.67). These results suggest that variation in RxA accounts for a substantial component of variation in the cortical and stele dimensions, but is less consistently associated with metaxylem traits.

Principal component analysis (PCA) placed roots along a continuous gradient from basal (N1) to more apical (N4) nodal positions (Figure 2). PC1 and PC2 explained 55.4% and 26.5% of variation, respectively. RxA and cortical traits contributed strongly to PC1, whereas stele traits contributed predominantly to PC2. N1 and N4 occupied opposite ends of the main gradient, with N2 and N3 in intermediate positions. Thus, nodal positions differed in their overall anatomical phenotype, although the distributions overlapped rather than forming discrete groups.

### 3.3. Nodal root anatomical trait trajectories are broadly maintained across growth environments

Several positional trends were reproduced in greenhouse-grown plants (Figure S2). RxA, CA, CCFN, SxA, and CSR generally increased from N1 to N4, whereas tMXA and MSR declined. However, the magnitude of cortical changes was smaller in the greenhouse than in the field. Metaxylem traits differed between environments. Mean metaxylem area declined in both environments (field slope = −0.21; greenhouse slope = −0.087; both *P <* 0.001), but this variation was associated with a decline in total metaxylem area in the field and an increase in vessel number in the greenhouse. Greenhouse PCA similarly separated RxA and cortical traits from vascular traits along the first two components (40.9% and 28.5% of variation, respectively). These results indicate broad consistency in the direction of positional variation, together with environmental differences in its magnitude and anatomical basis.

### 3.4. Anatomical traits show node-dependent allometric relationships

To determine how internal tissues scaled with RxA, we fitted allometric relationships between RxA and individual anatomical traits at each nodal position (Figure 3). Cortex and stele areas generally showed stronger RxA relationships than metaxylem traits. Fits for CCFN and mCCD were intermediate (R² ≈ 0.18 to 0.64 for field and 0.11 to 0.46 for the greenhouse dataset), whereas metaxylem traits (MXN, mMXA, and tMXA) frequently showed weak or nonsignificant relationships, consistent with the greenhouse dataset (R² ≤ 0.31). ANCOVA revealed significant interactions between RxA and nodal position, indicating that the relationship between RxA and individual anatomical traits was not constant across nodal positions on the main culm (Table S3, Figure S4). While specific exponent values varied slightly between the field and greenhouse, their developmental trends from N1 to N4 were consistent. Scaling exponents progressively decreased for cortical traits (CA, mCCD, and CCFN) across successive nodes, but increased for metaxylem area traits (tMXA and mMXA), with MXN remaining largely stable (Figure S5).

**Figure 3.**
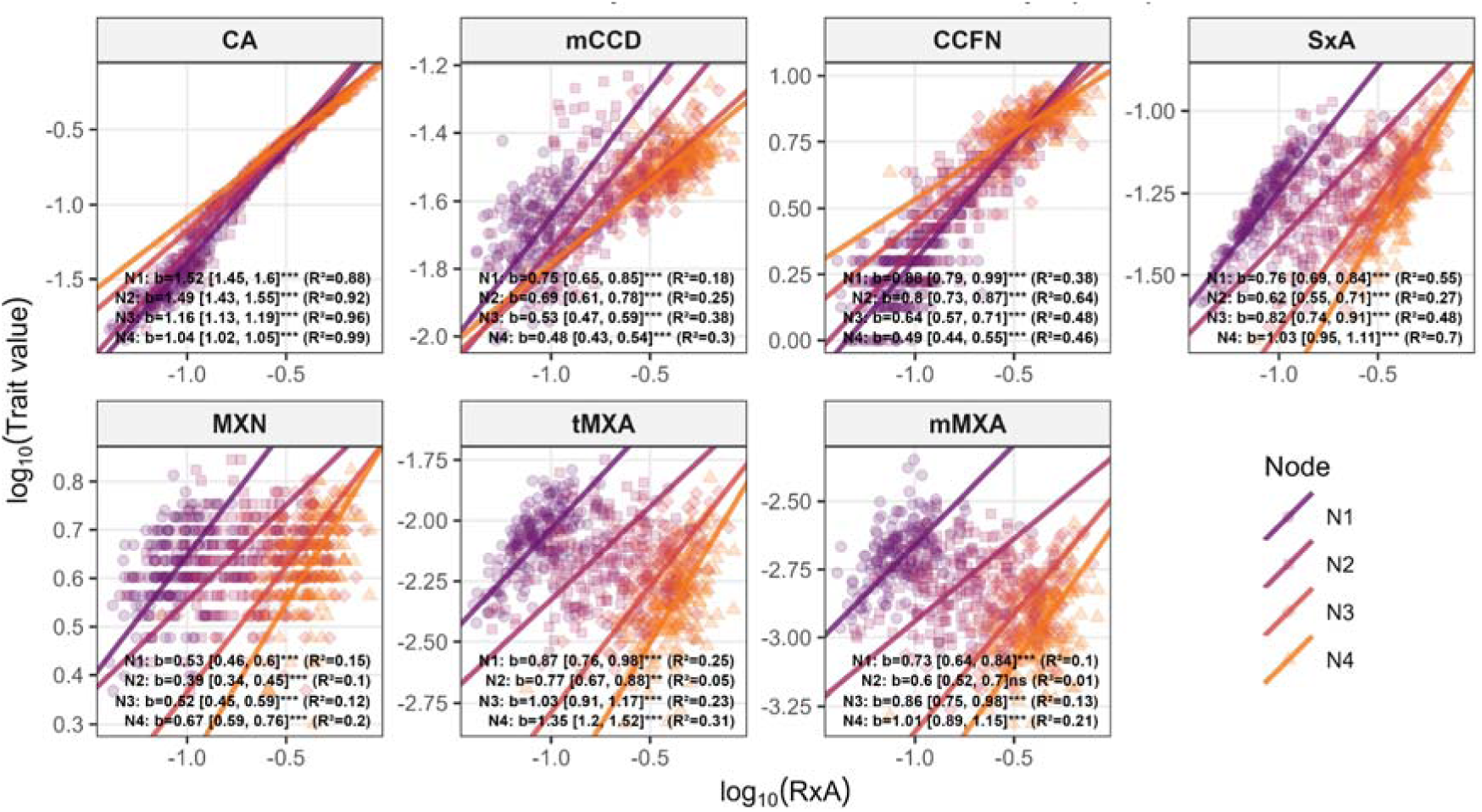
Node-specific allometric scaling relationships for the field dataset. Standardised Major Axis (SMA) regressions of log10-transformed trait values against log10-transformed root cross-sectional area (RxA) across nodal positions (N1–N4). Annotations detail the SMA scaling exponent (*b*, i.e., *b*_node_) with 95% CI, variance explained (R²), and association significance (* *P* < 0.05, ** *P* < 0.01, *** *P* < 0.001, ns = not significant) for each node-trait combination. Trait abbreviations: CA, cortex area; CCFN, cortical cell file layer number; mCCD, mean cortical cell diameter; SxA, stele cross-sectional area; MXN, metaxylem vessel number; tMXA, total metaxylem area; mMXA, mean metaxylem area.

### 3.5. Size correction reveals differences in tissue allocation among nodal positions

After standardisation to a common RxA (*x*_0,common_), several anatomical trends persisted, although their magnitude changed from the raw unadjusted trait values, where differences in GLMM slopes were notably large (Figure 4, Figure S9, Figure S10). This allometric correction reduced the dependence of anatomical traits on root cross-sectional area (RxA) (Figure S8, Figure S11, Figure S12). The increase in cortical area was substantially attenuated (Field: raw slope = 0.775; adjusted slope = 0.157; both *P <* 0.001). Similar attenuation was observed for CSR (Field: 0.707 to 0.483), CCFN (Field: 0.4 to 0.217), and MSR (Field: −0.261 to −0.184; all *P <* 0.001). Therefore, much of the raw variation in cortical traits was associated with increasing RxA from basal to apical nodal positions. Nevertheless, distinct positional differences remained at a common RxA.

**Figure 4.**
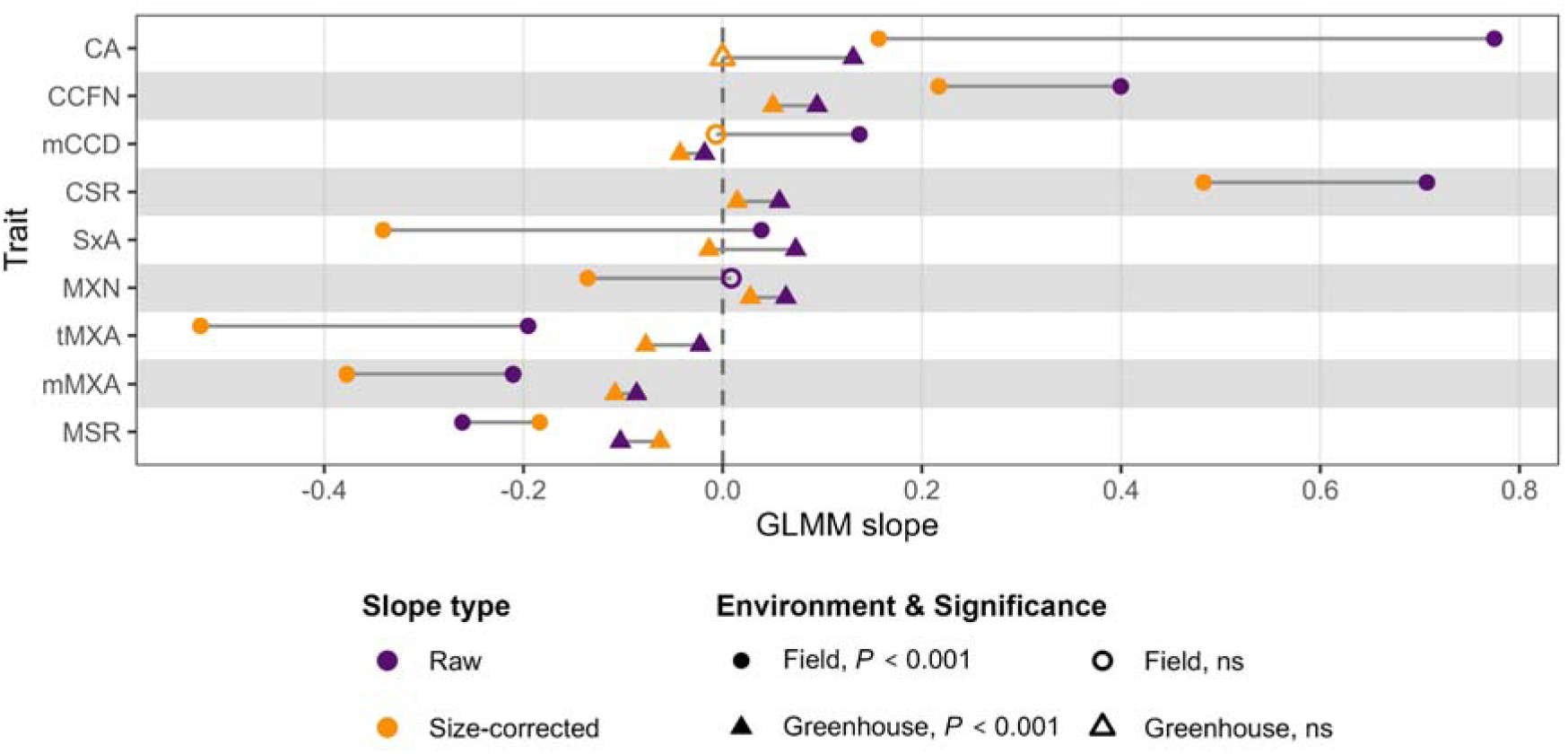
Generalised linear mixed model (GLMM) slopes represent the trajectories of root anatomical traits across nodes. The plot contrasts the slopes of trait gradients across successive developmental nodes (N1–N4) using raw (unadjusted) values versus size-corrected values in both field and greenhouse environments, highlighting the directional shift in slopes after removing the confounding effect of RxA. Trait abbreviations: CA, cortex area; CCFN, cortical cell file layer number; mCCD, mean cortical cell diameter; CSR, cortex-to-stele area ratio (CA/SxA); SxA, stele cross-sectional area; MXN, metaxylem vessel number; tMXA, total metaxylem area; mMXA, mean metaxylem area; MSR, metaxylem-to-stele area ratio (tMXA/SxA). Significance levels for the GLMM slopes: ns, not significant (*P* ≥ 0.05); *P* < 0.001.

The strongest change occurred in the stele area. SxA increased slightly across positions in the raw data (slope = 0.039, *P <* 0.001), but declined after size correction (slope = −0.341, *P <* 0.001) (Figure 4, Figure S9). This observation was also conserved in the greenhouse dataset (slope = -0.014, *P <* 0.001) (Figure 4, Figure S10). At a common RxA, roots from more apical nodes therefore had smaller steles than roots from basal nodes. Metaxylem traits showed similar trends after correction for RxA. MXN was not observed to be associated with nodal position in the raw data (Field: slope = 0.009, non-significant), but declined after adjustment (slope = −0.136, *P <* 0.001). Consequently, the negative gradients in tMXA and mMXA became much steeper after correction (−0.195 to −0.524 and −0.210 to −0.377, respectively; all *P <* 0.001).

Furthermore, the size correction also distinguished two distinct cellular components of cortical anatomy. While both CCFN and mCCD increased across nodal positions in the raw data, size correction indicated that the increase in mCCD was largely associated with variation in RxA; the size-corrected slope became non-significant in the field (slope = -0.007) and significantly negative in the greenhouse (slope = -0.043, *P <* 0.001). Conversely, CCFN continued to increase significantly across nodes in both environments (Field slope = 0.217; Greenhouse slope = 0.050; both *P <* 0.001). These results suggest that greater cortical cell file layer number, rather than increased mean cortical cell diameter, contributes more strongly to positional differences in cortical organisation at a common RxA (Figure 4).

The size-corrected trait correlations revealed negative associations between several cortical and stele traits across nodal positions (Figure S9, Figure S10). In the corresponding PCA biplot of the field dataset, PC1 explained 72.1% of the variation and opposed cortical traits (CA, CSR, CCFN) against vascular traits (tMXA, mMXA, SxA). PC2 explained 13.8% of the variance and was associated mainly with MXN and mCCD. Nodal root cohorts (N1–N4) remained largely separated along PC1, reflecting differences in cortical and vascular tissue organisation across successive nodal positions. Together, these analyses support a continuous difference in tissue allocation, with the clearest distinction between the most basal position and more apical positions.

Together, these allometric adjustments reveal differences in tissue allocation across successive nodal cohorts. At a common RxA (), roots emerging from more apical nodes exhibited smaller stele and metaxylem areas and relatively greater cortical allocation. The latter was associated primarily with increased cortical cell file layer number rather than mean cortical cell diameter. The higher percentage of variance explained by PC1 in the field (72.1%) compared with the greenhouse (47.7%) indicates that a greater proportion of anatomical variation was captured by the first principal component under field conditions.

### 3.6. Genetic associations differ between raw and size-corrected anatomical traits

Genotypic repeatability (*R*) for root anatomical traits varied significantly across environments, nodes, and between raw and size-corrected measurements. Greenhouse conditions yielded consistently higher *R* estimates across most traits than the field environment, where micro-environmental heterogeneity diluted the genetic signal, particularly at early nodes (N1 and N2). Metaxylem and stele-related traits, specifically total metaxylem area (tMXA), mean metaxylem area (mMXA), and the metaxylem-to-stele ratio (MSR), exhibited the highest and most stable repeatability, peaking at more apical nodal positions (N3 and N4) with greenhouse *R* values reaching 0.50 to 0.65. In contrast, cortical cell file layer number (CCFN) and mean cortical cell diameter (mCCD) showed consistently poor repeatability (*R* < 0.30) across all experiments. The application of size correction generally caused a marginal reduction in *R* for structural traits relative to raw values, confirming that a portion of the unadjusted genetic variance is driven by RxA. Nevertheless, the size-corrected data retained robust genetic signals for key tissue-partitioning traits, supporting their use in GWAS after accounting for variation in RxA (Figure S13).

Genetic variance estimates varied by trait, nodal position, and environment. Size correction generally retained broad differences among traits in the reported repeatability estimates, although some estimates changed considerably (Figure S13).

Genome-wide association analyses were conducted separately by trait, nodal position, and environment (Figure S14, Table S7 and Table S8). Using the reported suggestive threshold (−log_10_ *P* ≥ 3.64; FarmCPU), the raw-trait analyses identified 213 (164 unique between environments) associated SNPs in the field and 132 (123 unique between environments) in the greenhouse. In comparison, models utilising allometrically size-corrected traits detected 170 (162 unique) and 102 (79 unique) SNPs in the field and greenhouse, respectively. In the raw field analysis, associations were distributed across cortical traits, vascular traits, and RxA, with comparatively limited overlap among these categories. This limited intersection between cortical and vascular trait associations persisted even after size correction.

The detected genetic associations for nodal root anatomy were highly trait-and environment-specific, with limited overlap among traits (Figure 5, Figure S15). Under field conditions, associations were predominantly identified for RxA and cortical area traits, whereas greenhouse associations were more frequently identified for metaxylem-related traits (mMXA, tMXA), particularly at N4. Applying size corrections identified additional SNP associations for nearly all measured traits that were not detected using raw anatomical measurements. Although the sets of associated SNPs were largely distinct, some overlap occurred between raw and size-corrected versions of the same traits, suggesting that some associations were retained after accounting for variation in RxA. Few lead SNPs were shared among nodal positions or between field and greenhouse environments. Thus, the detected associations depended on trait, nodal position, and environment. However, limited overlap among suggestive associations does not necessarily indicate distinct genetic control, as the detection of associations also depends on allele effects, trait variation, linkage disequilibrium, and statistical power.

**Figure 5.**
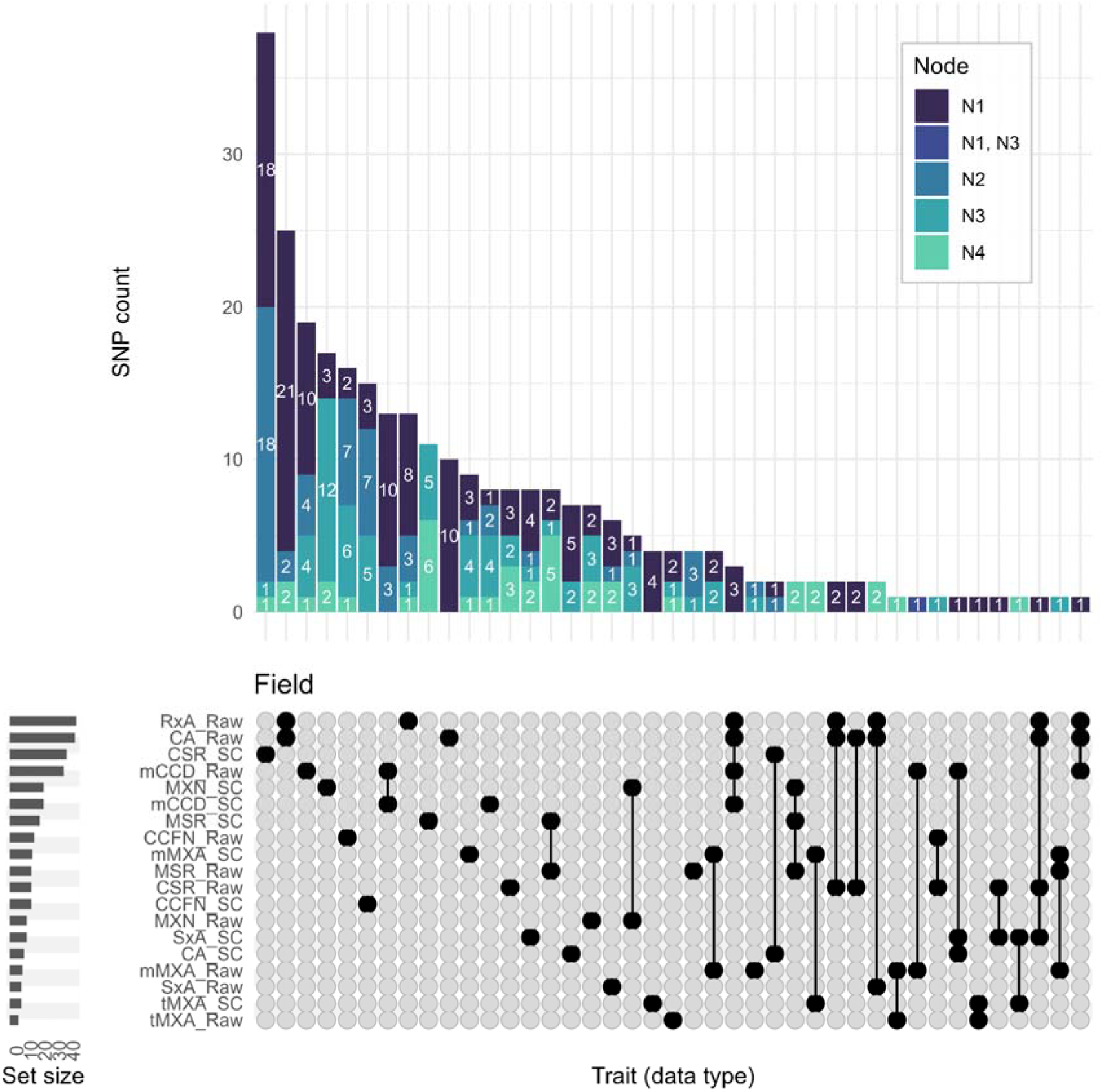
Shared and unique genetic loci associated with raw and size-corrected root anatomical traits within the field environment. UpSet plot illustrating the intersection of suggestive (−log10 P ≥ 3.64; FarmCPU model) single-nucleotide polymorphisms (SNPs) across unadjusted (“_Raw”) and size-corrected (“_SC”) traits. Vertical bars denote the number of SNPs unique to a specific trait-version combination or shared across multiple sets. Bar segments are coloured according to the specific nodal positions (N1–N4) from which the significant associations were mapped. Horizontal bars display the total number of suggestive SNPs detected for each individual trait version. Trait abbreviations: CA, cortex area; CCFN, cortical cell file layer number; mCCD, mean cortical cell diameter; CSR, cortex-to-stele area ratio (CA/SxA); SxA, stele cross-sectional area; MXN, metaxylem vessel number; tMXA, total metaxylem area; mMXA, mean metaxylem area; MSR, metaxylem-to-stele area ratio (tMXA/SxA).

### 3.7. Functional annotations differ between association sets

Candidate-gene sets assigned to raw and corrected associations partly overlapped, with 243 genes in the field and 169 in the greenhouse, as reported by the enrichment analysis (Figure 6). In the field, genes unique to the size-corrected analysis showed enrichment for functions associated with cell-wall processes (e.g., lignin and cellulose biosynthesis) and transport (e.g., carbohydrate and plasmodesmata-mediated transport). In contrast, genes from raw associations more often included growth- and signalling-related terms (e.g., ethylene-activated signalling). A keyword-based comparison of these functional categories was significant in the field (chi-square *P* = 0.026), but not in the greenhouse (chi-square *P* = 0.620) (Figure S16). These patterns suggest that trait adjustment changes the composition of candidate-gene sets; functional classifications remain exploratory and do not establish causal pathways.

**Figure 6.**
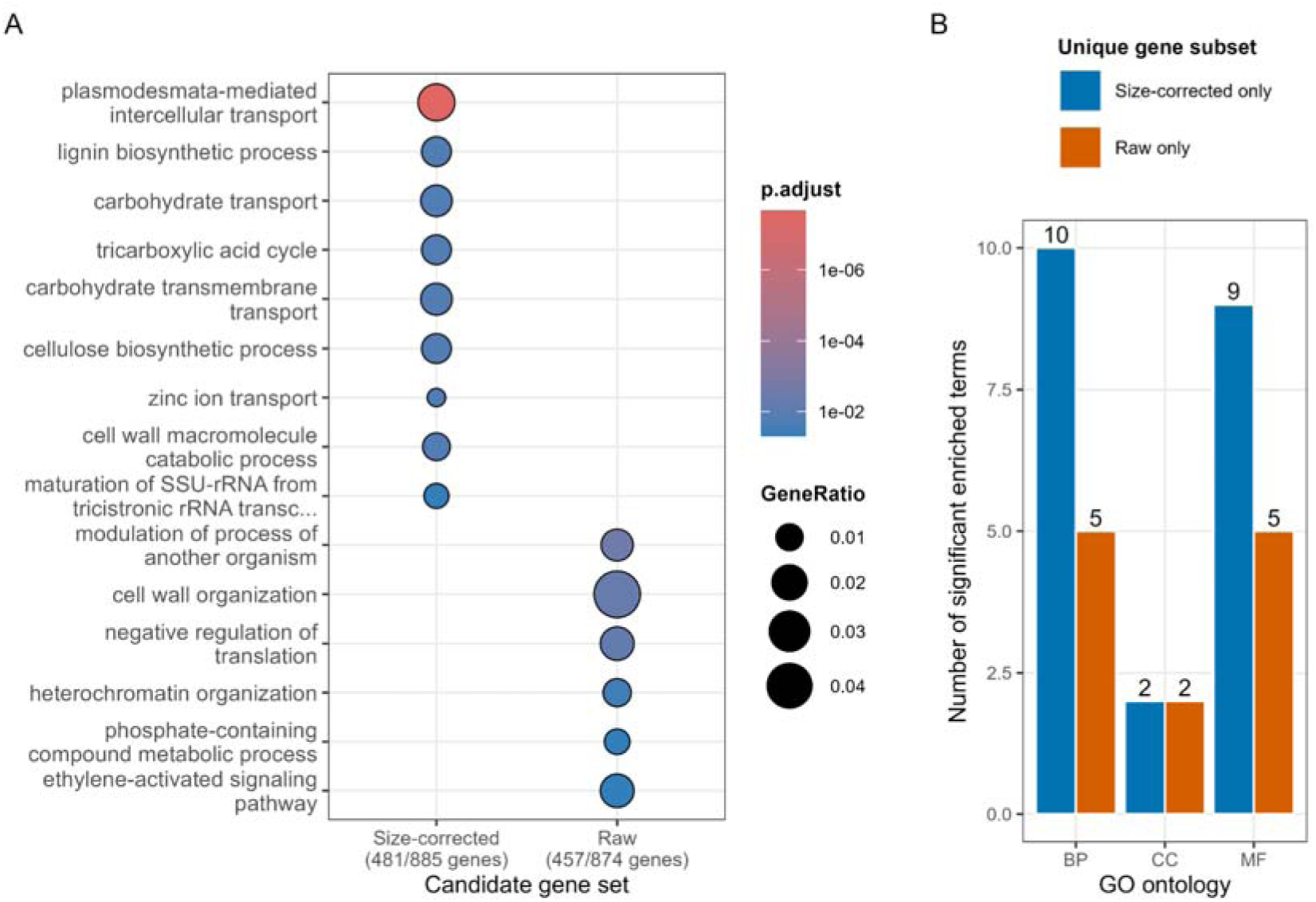
Functional enrichment and coherence of candidate genes under field conditions, where all nodal cohorts and traits are consolidated. (A) Side-by-side dot plot of significantly enriched Biological Process (BP) Gene Ontology terms for the size-corrected (481 genes) and unadjusted raw (457 genes) GWAS models. Node size represents the gene ratio, and colour indicates the Benjamini-Hochberg adjusted p-value. (B) Functional coherence of transformation-specific genes, illustrating the total number of significant GO terms (Biological Process [BP], Cellular Component [CC], and Molecular Function [MF]) derived from candidate genes uniquely identified by either the size-corrected or unadjusted models. The size-corrected unique gene set exhibits higher functional coherence, yielding nearly twice as many significant BP terms as the unadjusted unique set. Details about the greenhouse dataset are mentioned in Figure S16.

### 3.8 Representative genes from the raw and size-corrected data

Gene expression data revealed 76 root-expressed genes associated with allometrically size-adjusted traits and 64 root-expressed genes associated with raw, unadjusted root traits (Table S10, Table S11), within the core SNP confidence interval of ±200 Kb across environments and nodal root positions. Furthermore, we identified 32 genes shared between the allometric and raw unadjusted trait sets, specifically associated with mCCD, mMXA, MSR, and MXN. Interestingly, integrating GWAS associations with publicly available gene expression data identified highly root-expressed candidate genes associated with anatomical traits across different nodal positions, yielding partially distinct candidate gene sets from raw and size-corrected analyses.

Together, GWAS association studies suggest convergent and distinct regulation of RxA-independent and RxA-dependent root anatomical traits at different root node levels, a pattern reported for the first time in cereals, specifically wheat.

## 4. Discussion

### 4.1. Successive nodal roots form a developmentally structured root system

Cereal root systems are assembled progressively throughout plant development. Unlike embryonic roots, which emerge during germination and the seedling stage, nodal roots emerge sequentially from successive culm nodes as the plant develops (*6*, *7*). This organisation results in a mature root system composed of roots initiated at different stages of whole-plant development. The contribution of nodal roots to the overall root system also increases with plant age (*75*, *6*, *76*, *17*). Thus, the progressive emergence of nodal roots changes not only the composition of the root system but also the relative contribution of different root classes throughout plant development.

Our results demonstrate that this developmental sequence is accompanied by systematic changes in the anatomy of the roots produced. Roots emerging from successive main culm nodes differed in RxA, cortical and stelar dimensions, and metaxylem anatomy. Root cross-sectional area and cortical allocation increased from basal to more apical nodal positions, whereas total and mean metaxylem area declined. Multivariate analyses similarly revealed a continuous gradient of anatomical variation across successive nodal positions rather than discrete anatomical groups. These findings extend observations in maize, where roots originating from successive stem nodes differ in anatomical and architectural traits (*34*, *35*, *37*), and indicate that anatomical differentiation within the post-embryonic root system is also an important feature of wheat (*8*, *25*, *77*).

An important distinction, however, is that nodal position is associated with several developmental variables. Roots originating from basal nodes emerge earlier and are consequently older at a common harvest stage, whereas roots from more apical nodes emerge later and develop under a different whole-plant physiological context (*6*, *78*). Nodal position, root age, emergence time, and plant developmental stage are therefore not independent in the present experiment. The observed anatomical gradients may consequently reflect differences in root developmental age, the physiological conditions under which roots develop, or position-associated developmental regulation and RxA (*34*, *38*).

This organisation parallels the sequential development of aboveground organs. Cereal shoots are composed of successive phytomers, with leaves and internodes at different positions exhibiting distinct morphological and physiological characteristics (*6*, *79–81*). Extending this developmental framework belowground suggests that nodal roots develop in structural coordination with their corresponding parent phytomer (*10*). Consistent with this interpretation, we observed a positive relationship between stem diameter and root cross-sectional area (R² = 0.35, *P <* 0.001; Figure S17), indicating that the dimensions of nodal roots are partly associated with those of the main culm node (*6*, *82*). However, considerable variation in RxA remained unexplained by stem diameter. Furthermore, roots from more apical positions were larger than basal roots even within overlapping ranges of stem diameter. These observations suggest that anatomical differences among nodal positions cannot be attributed solely to differences in parent culm size. Because roots from successive nodal positions emerge at different times, they also differ in age and developmental history. Therefore, the anatomical differences observed here cannot be attributed exclusively to nodal position, but may also reflect differences in root age, whole-plant developmental stage, and the environmental conditions experienced during root development.

Together, these findings support a view of the wheat root system as a developmentally organised series of anatomically distinct nodal roots. Rather than treating all post-embryonic nodal roots as equivalent, their position within the developmental sequence should be considered when characterising root phenotypes and interpreting their potential functions (*34*, *37*).

### 4.2. Allometric relationships reveal differences in tissue organisation among nodal positions

A major challenge in interpreting anatomical variation among nodal roots is distinguishing changes in tissue organisation from differences associated with RxA (*44*, *83*). Larger roots generally contain more cortical and vascular tissue, and the dimensions of individual tissues often scale with root cross-sectional area (*45*, *83*, *84*). Consequently, anatomical differences among successive nodal positions could arise simply because later-emerging roots are larger. This distinction is particularly important when comparing tissue dimensions across root classes or developmental stages (*34*, *85*).

Our allometric analysis showed that root cross-sectional area (RxA) accounted for a substantial component of anatomical variation, particularly in cortical traits. The strong increase in cortical area from N1 to N4 was considerably attenuated after size correction, indicating that much of the observed cortical expansion was associated with increasing RxA. However, several anatomical gradients persisted after standardisation to a common RxA. In particular, the cortex-to-stele area ratio remained higher in roots from more apical nodes, whereas stele area and metaxylem traits declined. These findings demonstrate that anatomical variation across successive nodal positions cannot be explained by differences in RxA alone (Figure 4, Figure S9, Figure S10).

The most pronounced change occurred in the stele area. Although the stele area increased slightly from basal to apical nodes in the raw data, this relationship was reversed after size correction (Figure 4). At a common RxA, roots originating from more apical nodes had smaller steles than roots from basal nodes. Because total RxA constrains the space available for individual tissues, greater cortical allocation necessarily corresponds to a smaller proportion of the cross-section occupied by the stele. However, whether this reflects differences in the developmental regulation of cortical and vascular tissues, or constraints on their relative expansion, remains unclear. These results indicate that the relative organisation of cortical and vascular tissues varies across successive nodal positions, even after accounting for differences in RxA (*83*, *84*).

Size correction also revealed differences in the cellular basis of cortical expansion. Both cortical cell file number and mean cortical cell diameter increased across successive nodes in the raw data. However, after accounting for RxA, the positional relationship with mean cortical cell diameter was no longer significant, whereas cortical cell file number remained positively associated with nodal position. Thus, at a comparable RxA, roots from more apical nodes generally contained more cortical cell layers without a corresponding increase in mean cortical cell diameter. This suggests that differences in cortical organisation are associated more strongly with cortical cell file number than with radial enlargement of individual cortical cells (*34*, *37*, *86*). This distinction may have functional implications, as cortical cell size and cell file number can influence the metabolic costs of root construction and maintenance (*23*, *30*). Larger cortical cells and fewer cortical cell files have been associated with reduced metabolic costs, whereas additional cortical cell layers may contribute to root mechanical properties and radial transport. Thus, the greater cortical cell file number observed in roots from more apical nodes could reflect different functional requirements, although these consequences were not directly evaluated in this study.

Importantly, the allometric relationships themselves also varied among nodal positions. Cortical area showed a stronger increase with RxA in basal roots, whereas the scaling exponents of several metaxylem traits were greater in roots from more apical nodes. This indicates that the relationship between overall RxA and individual tissue dimensions is not constant across the developmental sequence. Consequently, anatomical differentiation among nodal positions involves both changes in absolute tissue dimensions and differences in how tissues scale with RxA.

These patterns have potential functional implications. Cortical traits influence the metabolic costs of root construction and maintenance, as well as radial transport and mechanical properties, while stele and metaxylem dimensions are important determinants of hydraulic transport (*25*, *45*, *86*). The greater cortical allocation and reduced vascular dimensions in roots emerging from more apical nodes at a common RxA could therefore influence the balance between metabolic costs, mechanical properties, and water transport capacity. In particular, differences in metaxylem vessel dimensions may influence axial hydraulic conductance, which depends strongly on vessel radius. However, anatomical variation alone does not establish functional differences. Future studies integrating anatomical allometry with measurements of root respiration and theoretical axial hydraulic conductance, alongside direct measurements of water and nutrient uptake, could help determine the functional consequences of these anatomical differences.

### 4.3. Anatomical differentiation is broadly maintained across growth environments

The general trends in anatomical traits across successive nodal positions were broadly consistent between field and greenhouse conditions. Root cross-sectional area, cortical area, cortical cell file layer number, stele area, and cortex-to-stele ratio generally increased toward more apical nodes, whereas the proportion of stele area occupied by metaxylem area declined. Although the absolute trait values differed between environments, the broadly similar trends from N1 to N4 suggest that positional variation is a recurring feature of wheat nodal root development.

However, the magnitude of anatomical variation across nodal positions differed between environments, particularly for cortical and metaxylem traits. Changes in cortical dimensions across nodal positions were generally less pronounced in greenhouse-grown plants than in field-grown plants (Figure 2, Figure S2). Although mean metaxylem area declined across nodal positions in both environments, the associated changes in total metaxylem area and vessel number differed. Total metaxylem area declined more strongly in the field, whereas metaxylem vessel number remained relatively stable in the field but declined in the greenhouse. Thus, similar positional trends in individual traits can be accompanied by different patterns of anatomical variation between environments.

These differences may reflect the environmental conditions experienced during root development. Successive nodal roots are produced at different times and may develop under varying soil strength, temperature, water availability, and nutrient supply. Previous studies in maize have similarly demonstrated that anatomical variation among nodal positions can interact with environmental conditions, including nutrient and water availability (*34*, *35*). Our results suggest that anatomical trends across successive nodal positions are broadly consistent between environments, although the magnitude of these trends and the underlying anatomical characteristics can differ.

This variation also has implications for the interpretation of genotype-by-environment interactions. Genetic differences in root anatomy may not be expressed uniformly across all nodal positions. Instead, genotypes may differ in their overall anatomical phenotype, the magnitude of positional variation, or the scaling relationships between individual tissues and RxA (*34*, *87*). Characterising these relationships across multiple nodal positions and environments may therefore provide a more complete description of anatomical variation than measurements from a single root position.

### 4.4. Integrating allometry into root phenotyping reveals distinct genetic associations

From a phenotyping perspective, these results emphasise the importance of recording nodal position and accounting for variation in RxA when comparing anatomical traits (*37*, *88*). Measurements from a single nodal position may not represent the anatomical characteristics of the mature root system, while pooling roots from different positions may obscure systematic developmental variation (*34*). When comparing plant anatomical or architectural traits, differences in overall size, such as length, area, volume, or biomass, can distort the data. Allometric analysis allows anatomical traits to be compared after accounting for variation in RxA. Integrating this approach with information on nodal position provides a framework for identifying anatomical differences that are not explained by RxA alone. This may also help distinguish genetic associations related to overall RxA from those associated with tissue organisation.

Genetic effects on root dimensions can contribute indirectly to variation in cortical and vascular traits (*45*, *83*). Allometric correction provides a means of examining anatomical variation after accounting for relationships with RxA (*44*, *89*). Our parallel GWAS analyses identified largely different associations for raw and size-corrected anatomical traits, with limited overlap among nodal positions and between growth environments. This indicates that the detected genetic associations depend on how anatomical phenotypes are defined and the developmental position at which they are measured (*34*, *85*). These differences demonstrate that accounting for RxA can substantially alter which genomic regions are associated with anatomical variation.

The functional annotations of candidate genes provided additional differences between the two approaches. In field-grown plants, candidate genes identified exclusively by size-corrected analysis were enriched for functions associated with cell-wall processes and transport, whereas raw trait associations more frequently included growth- and stress-related functional categories. However, this distinction was less pronounced in greenhouse-grown plants. These results suggest that size adjustment can influence the functional composition of candidate-gene sets, although the biological significance of these differences remains uncertain. Gene Ontology enrichment provides information about functional associations among candidate genes but does not establish their causal involvement in anatomical development (*90*, *91*).

Among the identified loci, TRAESCS5A02G388300 was highly expressed in the roots and associated with stele-related root traits, specifically in raw unadjusted MXN and allometrically adjusted tMXA in nodes 3 and 2, respectively. This gene is orthologous to the previously described AtBGLC1 in *Arabidopsis thaliana (AT5G20950)*, which is expressed in both the primary and lateral root caps and regulates xyloglucan turnover, thereby promoting primary cell-wall remodelling (*92*, *93*). TRAESCS6D02G406000 was identified as a highly root-expressed, unique candidate associated with raw unadjusted mCCD in node 1. TRAESCS6D02G406000 is described as CALS2/GSL3 in *Arabidopsis thaliana (AT2G31960),* which belongs to a 12-member family of CALLOSE SYNTHASE (CALS)/GLUCAN SYNTHASE-LIKE (GSL) genes responsible for the biosynthesis of callose (β-1,3-glucan). Its paralog GSL8 was proposed to interact with PDLP5 to regulate callose deposition at plasmodesmata (*94*). TRAESCS2B02G236500 was detected as highly root-expressed and was a unique candidate associated with CCFN in node 3. TRAESCS2B02G236500 ortholog gene in barley, HvCslf3 (HORVU2Hr1G042350), is part of the Poaceae-specific CslF gene family responsible for the (1,4)-β-linked glucoxylan, which forms the backbone of the hemicellulose in the primary cell wall. Interestingly, it was reported that mutated CslF3 RNAi alleles in barley show reduced root length growth, a significant reduction in root cortical area, and a trend toward reduced cortical cell file number compared to the wild type (*95*).

The limited overlap among GWAS associations across nodal positions may also reflect differences in the genetic regulation of root anatomy during plant development. However, this interpretation requires caution. Differences in association detection can arise from variation in trait distributions, genetic variance, environmental effects, and statistical power (*91*). Similarly, loci detected using size-corrected traits cannot automatically be interpreted as controlling tissue patterning independently of overall root growth. Size correction removes the modelled relationship with the root cross-sectional area but does not establish the underlying genetic mechanism (*44*, *89*). Functional validation and additional genetic analyses will therefore be necessary to determine whether the associations identified here represent distinct regulators of tissue organisation.

Together, these findings demonstrate the value of integrating nodal position and allometric relationships into root anatomical phenotyping. Rather than treating nodal roots as equivalent sampling units or relying exclusively on raw anatomical measurements, this approach allows anatomical variation to be evaluated within its developmental context and relative to a defined reference RxA. Combining raw and size-corrected traits with genetic analyses provides a more comprehensive framework for investigating the genetic architecture of root anatomy and identifying anatomical variation that may otherwise be obscured by differences in overall root cross-sectional area.

## 5. Conclusions

Successive wheat nodal roots differ systematically in their anatomy, and these differences cannot be explained by variation in RxA alone. Although increasing root cross-sectional area accounts for much of the cortical expansion observed across successive nodal positions, differences in cortical cell file number, stele area, and metaxylem anatomy persist after allometric correction. Furthermore, the scaling relationships between RxA and individual tissues vary among nodal positions, indicating that anatomical differentiation involves changes in both tissue dimensions and their relative organisation.

These patterns are broadly maintained across greenhouse and field environments, although their magnitude and anatomical basis differ. The distinct genetic associations identified for raw and size-corrected traits further underscore the importance of accounting for allometry when characterising the genetic architecture of root anatomy, as well as when investigating the underlying developmental mechanisms and true physiological function of root traits. Together, our findings support a view of the mature wheat root system as a developmentally organised series of anatomically differentiated nodal roots rather than a collection of equivalent root structures. Understanding how these anatomical differences influence resource acquisition and root function will be important for identifying developmentally relevant root traits for crop improvement.

## Author contributions

ABT and HMS conceived the study. ABT, MS, and HMS designed the study. Plant growth and wet-lab work were performed by ABT, MS, ILV, DHJ, AW, DS, and PL. Dry-lab methodology was performed by ABT, AW, and PL. ABT, HMS, MS, and RF performed data analyses. Funding was acquired by HMS, AD, MA and MS. ABT, MS, and HMS drafted the manuscript. The wheat diversity panel was generated and made available by MS, JD, PC, and AB. AD, MA, and GG advised on conceptualisation. All authors contributed substantially to the revisions.

## Declaration of generative AI and AI-assisted technologies in the manuscript preparation process

During the preparation of the work, the authors used Claude Sonnet 5 (Anthropic), via the ZGPK platform (IPK Gatersleben), to assist with English phrasing and R code editing. The authors reviewed all AI-assisted content and take full responsibility for the final article.

## Code and data availability

Data are available in the article’s supplementary material.

## Acknowledgments

We thank Corinna Trautewig, Avani Vallayil, Darshan Damodaran, Pauline Hampe, Maria Schoen, Zhongyang Wang, Luisa Meier, Luisa Wolfgramm, Rashmi Aryal, Osalumen Philemon Irusota, Wirassaya Tobdan, Gopika Shaju, and Misbah Chaudhry for their assistance with plant evaluations and sample collections from the field and greenhouse. We thank Peter Schreiber, Enk Geyer, Gerald Fiedler, Oliver Schmidt, Stefanie Fischer, Kerstin Jacobs, Anke Mueller, and Kathrin Tiemann for their assistance with greenhouse and field experiment management.

## Abbreviations

Nodal roots NR: Main culm nodal (N) root positions (basal to apical) (Node 1 to Node 4; N1 to N4
GWAS: genome-wide association studies
RxA: Root cross-sectional area
SxA: Stele cross-sectional area
CA: Cortex area
CSR: Cortex to stele ratio
CCFN: Cortical cell file layer number
mCCD: Mean cortical cell diameter
MXN: Metaxylem vessel number
tMXA: Total metaxylem area
mMXA: Mean metaxylem area
MSR: Metaxylem to stele ratio

