## Supplemental Information for "Allometric analysis reveals distinct anatomical patterning and genetic architecture across wheat nodal roots"

### **Supplementary tables**

###### **Table S2. Summary table of the actual dataset obtained and the theoretical expected values mentioned in the top row.**

| Conditions | Node | Genotypes (n=190) | Average number of replicates (= total number of images/genotypes) (n=3) | Number of images obtained (n=570) |
| --- | --- | --- | --- | --- |
| Field | N1 | 190 | 2.69 | 511 |
| Field | N2 | 190 | 2.89 | 549 |
| Field | N3 | 190 | 2.93 | 557 |
| Field | N4 | 190 | 2.98 | 566 |
| Greenhouse | N1 | 190 | 2.92 | 555 |
| Greenhouse | N2 | 190 | 2.86 | 544 |
| Greenhouse | N3 | 190 | 2.75 | 522 |
| Greenhouse | N4 | 185 | 2.37 | 438 |
| - | - |  |  | Total: 4242/4560 |

**Table S3. Trait-specific evaluation of allometric scaling slope heterogeneity across nodes.** Analysis of Covariance (ANCOVA) results testing the interaction term between log-transformed root cross-sectional area (log_10_ RxA) and nodal position (N1–N4) for each continuous anatomical trait. A non-significant interaction (*P* ≥ 0.05) indicates homogeneous slopes, confirming that a shared allometric scaling exponent can be applied across all root nodes in the given environment.

| conditions | anatomy_trait | slope_p | slopes_equal |
| --- | --- | --- | --- |
| Field | CA | 3.95E-27 | FALSE |
| Field | CCFN | 9.32E-08 | FALSE |
| Field | mCCD | 0.587215 | TRUE |
| Field | CSR | 1.48E-12 | FALSE |
| Field | SxA | 1.83E-17 | FALSE |
| Field | MXN | 0.007516 | FALSE |
| Field | tMXA | 4.44E-09 | FALSE |
| Field | mMXA | 3.62E-06 | FALSE |
| Field | MSR | 0.418552 | TRUE |
| Greenhouse | CA | 0.006536 | FALSE |
| Greenhouse | CCFN | 0.887599 | TRUE |
| Greenhouse | mCCD | 0.401864 | TRUE |
| Greenhouse | CSR | 0.051128 | TRUE |
| Greenhouse | SxA | 0.136863 | TRUE |
| Greenhouse | MXN | 0.72413 | TRUE |
| Greenhouse | tMXA | 0.854189 | TRUE |
| Greenhouse | mMXA | 0.643204 | TRUE |
| Greenhouse | MSR | 0.711336 | TRUE |

### **Supplementary figures**

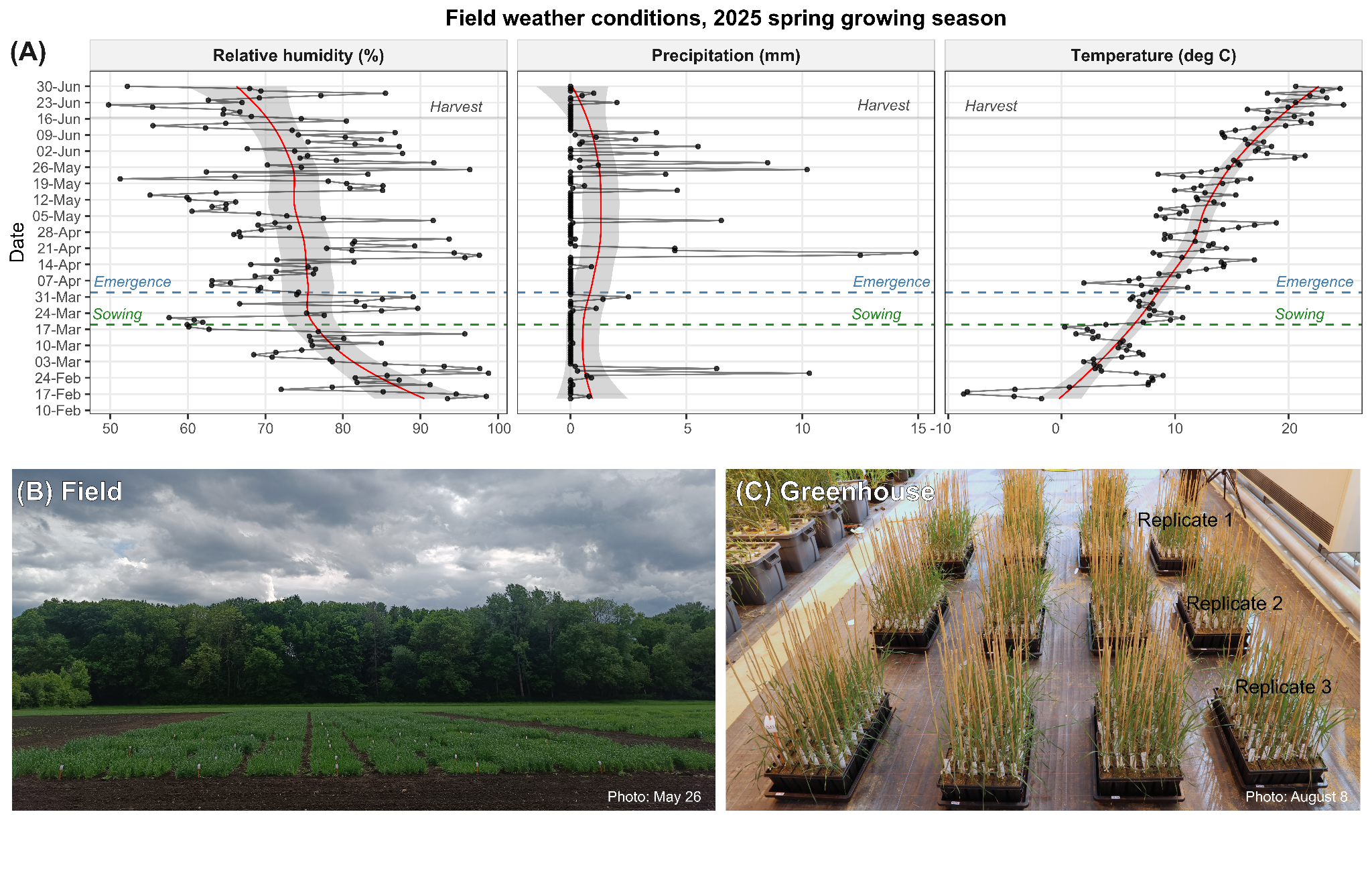

###### **Figure S1. Growth conditions for the wheat diversity panel. (A) Environmental conditions during the 2025 spring wheat growing season on the experimental field site (B) at Gatersleben, Germany.** Black dots represent daily measurements, while the red line with shading indicates the mean trend over the period. (C) Wheat diversity panel grown in quickpot trays under greenhouse-controlled conditions with randomised three replicate blocks (Replicate 1, 2 and 3). Key phenological dates: Field - sowing (March 19), emergence (April 2), and root harvesting at booting stage (June 16 - 17); Greenhouse - sowing (July 4), emergence (July 9), and root harvesting at booting stage (August 28).

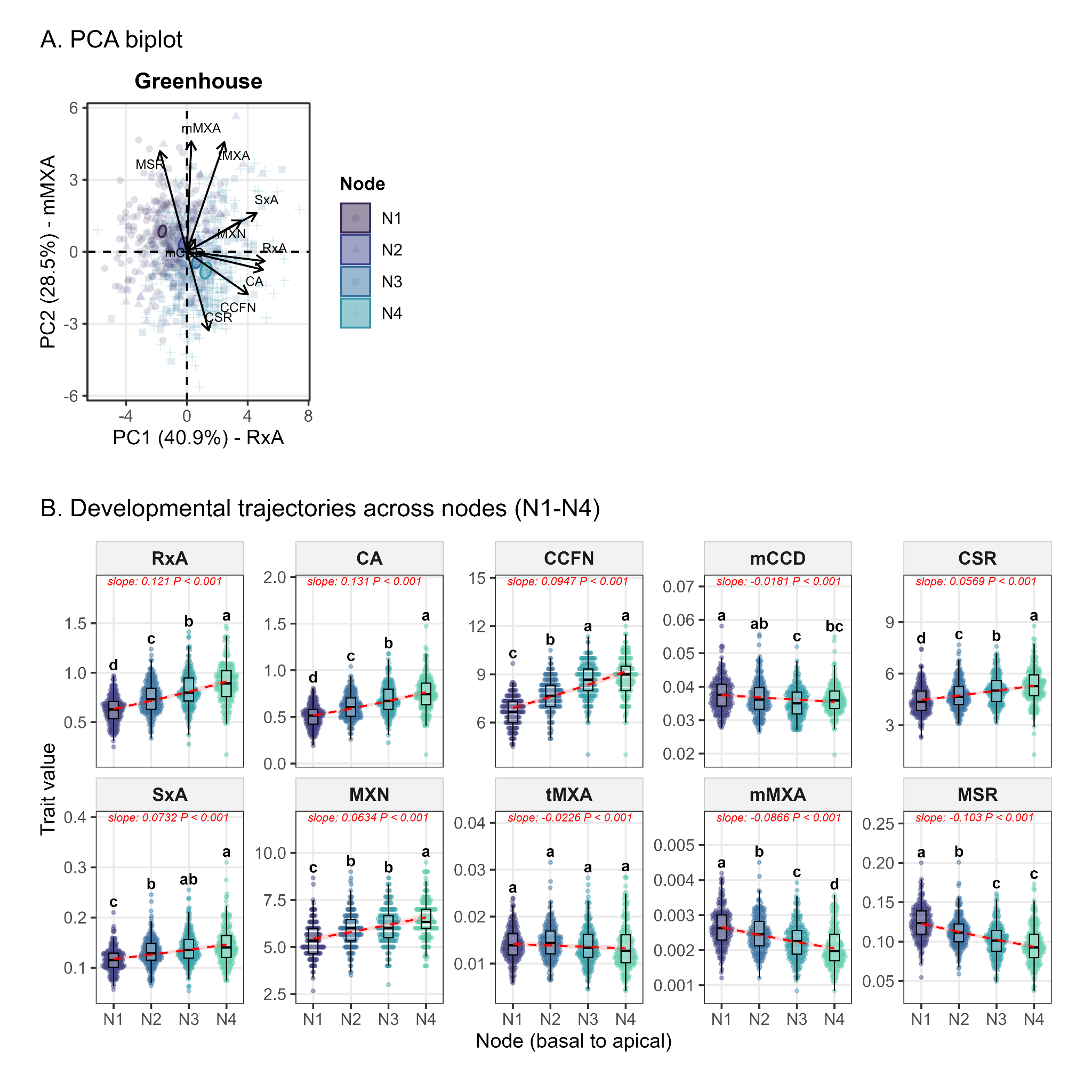

###### **Figure S2. Anatomical variation across successive nodal positions in greenhouse-grown wheat. Root anatomical traits vary systematically among roots emerging from four successive nodes (N1–N4) along the main culm, from basal to apical positions.** (A) Principal component analysis (PCA) of anatomical traits across four nodal positions. Genotype means were centred and estimated before analysis. Points are colored by nodal position, and ellipses represent 95% confidence regions. Arrows indicate trait loading vectors, and axes show the percentage of variance explained by each principal component. (B) Variation in root anatomical traits across nodal positions, including root cross-sectional area (RxA), cortex area (CA), cortical cell file layer number (CCFN), mean cortical cell diameter (mCCD), cortex-to-stele area ratio (CSR = CA/SxA), stele cross-sectional area (SxA), metaxylem vessel number (MXN), total metaxylem area (tMXA), mean metaxylem area (mMXA), and metaxylem-to-stele area ratio (MSR = tMXA/SxA). Boxplots show the distribution of genotype means at each nodal position. Dashed red lines and shaded bands indicate predicted trends and 95% confidence intervals from generalised linear mixed-effects models (GLMMs) fitted to replicate-level data, with nodal position as a fixed effect and genotype as a random intercept. Gamma models with a log link were used for continuous traits, Poisson models with a log link for count traits, and a beta model with a logit link for MSR. Model slopes and associated P-values are shown in red. Different lowercase letters indicate significant differences among nodal positions (Tukey's HSD, *P* < 0.05). The data represent 190 wheat genotypes, with each point corresponding to the mean of three biological replicates.

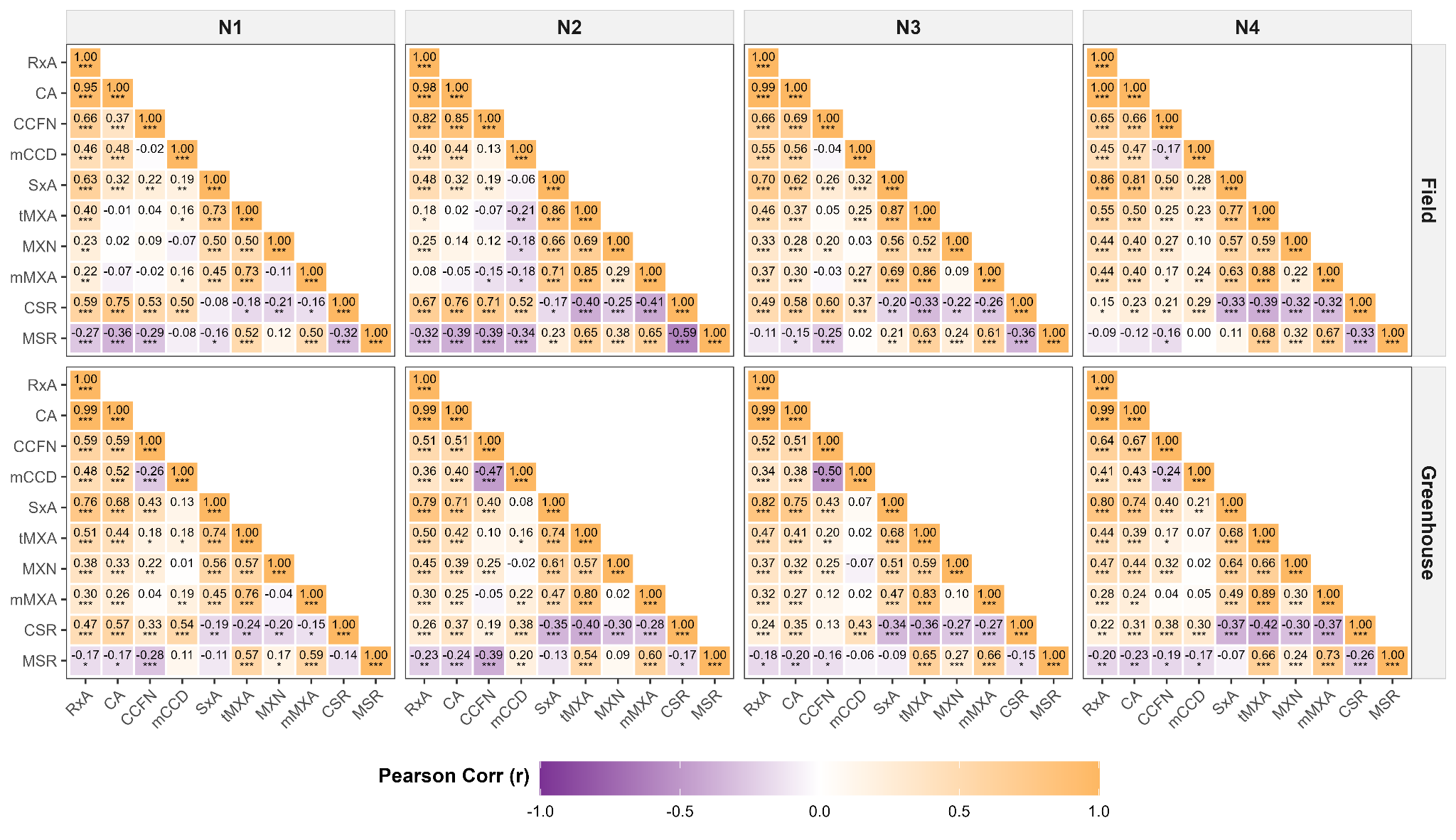

###### **Figure S3. Pairwise Pearson correlations between raw root anatomical traits across distinct nodal positions and environments.** The lower-triangle correlation matrices are faceted by root node (N1–N4, columns) and growth environment (Field vs. Greenhouse, rows). Statistical significance is denoted by asterisks stacked below the r values (*** *P* < 0.001, ** *P* < 0.01, * *P* < 0.05), and non-significant correlations (*P* ≥ 0.05) are displayed without asterisks. Trait abbreviations: CA, cortex area; CCFN, cortical cell file layer number; mCCD, mean cortical cell diameter; CSR, cortex-to-stele area ratio (CA/SxA); SxA, stele cross-sectional area; MXN, metaxylem vessel number; tMXA, total metaxylem area; mMXA, mean metaxylem area; MSR, metaxylem-to-stele area ratio (tMXA/SxA).

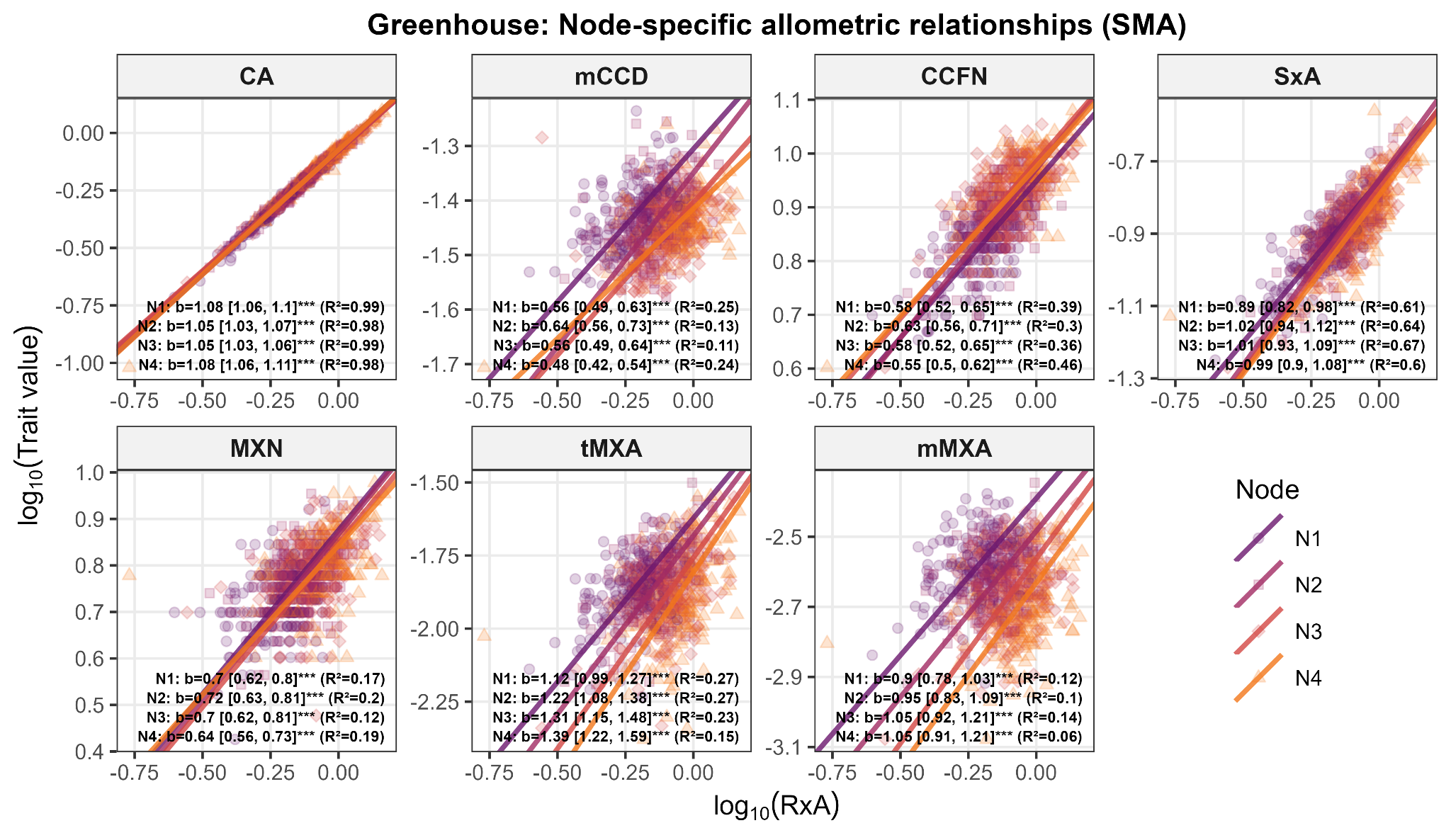

###### **Figure S4. Node-specific allometric scaling relationships for the greenhouse dataset. Standardised Major Axis (SMA) regressions of log10-transformed trait values against log10-transformed root cross-sectional area (RxA) across nodal positions (N1–N4).** Annotations detail the SMA scaling exponent (*b*) with 95% CI, variance explained (R²), and association significance (* *P* < 0.05, ** *P* < 0.01, *** *P* < 0.001, ns = not significant) for each node-trait combination. Trait abbreviations: CA, cortex area; CCFN, cortical cell file layer number; mCCD, mean cortical cell diameter; SxA, stele cross-sectional area; MXN, metaxylem vessel number; tMXA, total metaxylem area; mMXA, mean metaxylem area.

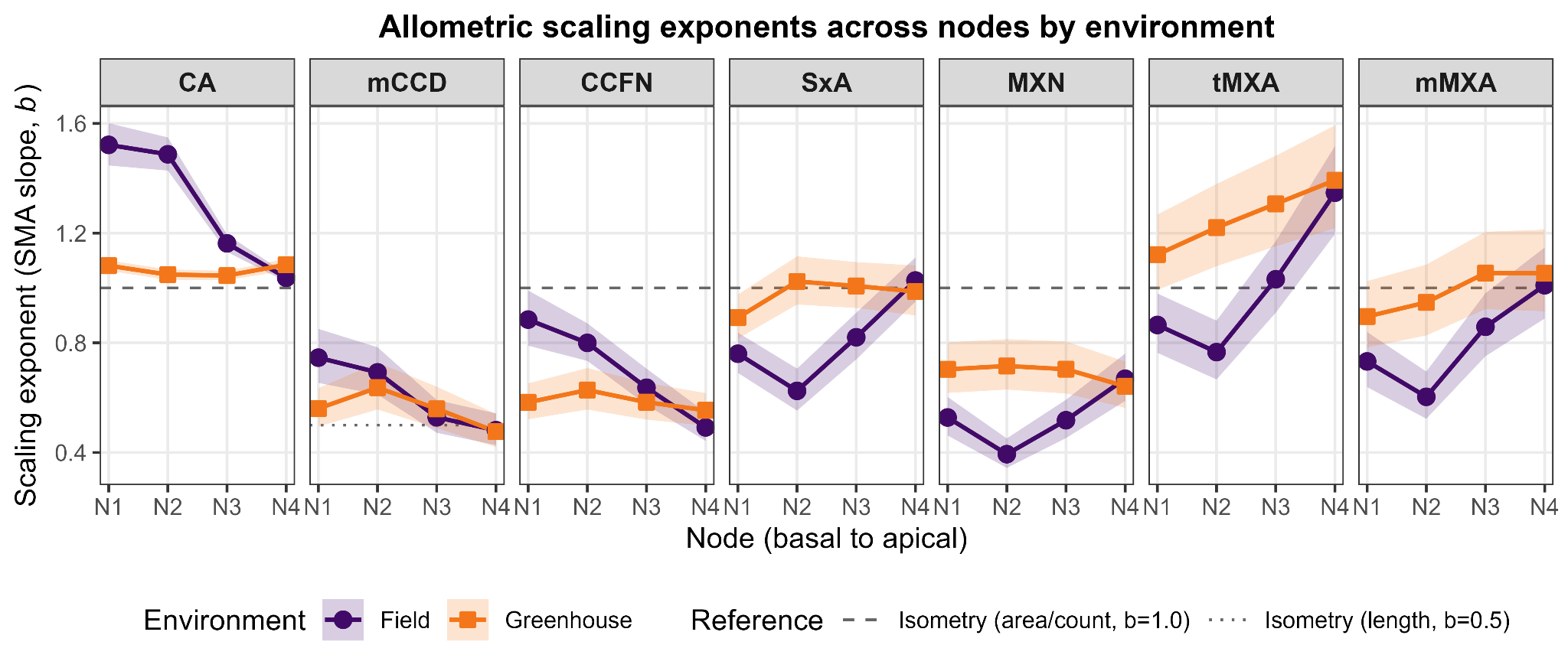

###### **Figure S5. Across different nodal positions under field- and greenhouse-grown wheat plants, allometric scaling exponents (*b*) were determined for root anatomical traits.** These *b* values represent the slopes derived from raw trait data subjected to standard major-axis regression, with the transformation log_10_(trait) ~ log_10_(RxA), as shown in Figure S4. The lines connecting symbols represent the traits for which the variation in b values is shown, and the shaded ribbon represents the 95% CI for the SMA slope estimate. The expected isometric values (*b_iso_*) vary with trait dimensionality: *b_iso_* = 1.0 for meristic, density, and area traits; and *b_iso_* = 0.5 for linear/length traits; denoted by dashed line types. Trait abbreviations: CA, cortex area; CCFN, cortical cell file layer number; mCCD, mean cortical cell diameter; SxA, stele cross-sectional area; MXN, metaxylem vessel number; tMXA, total metaxylem area; mMXA, mean metaxylem area.

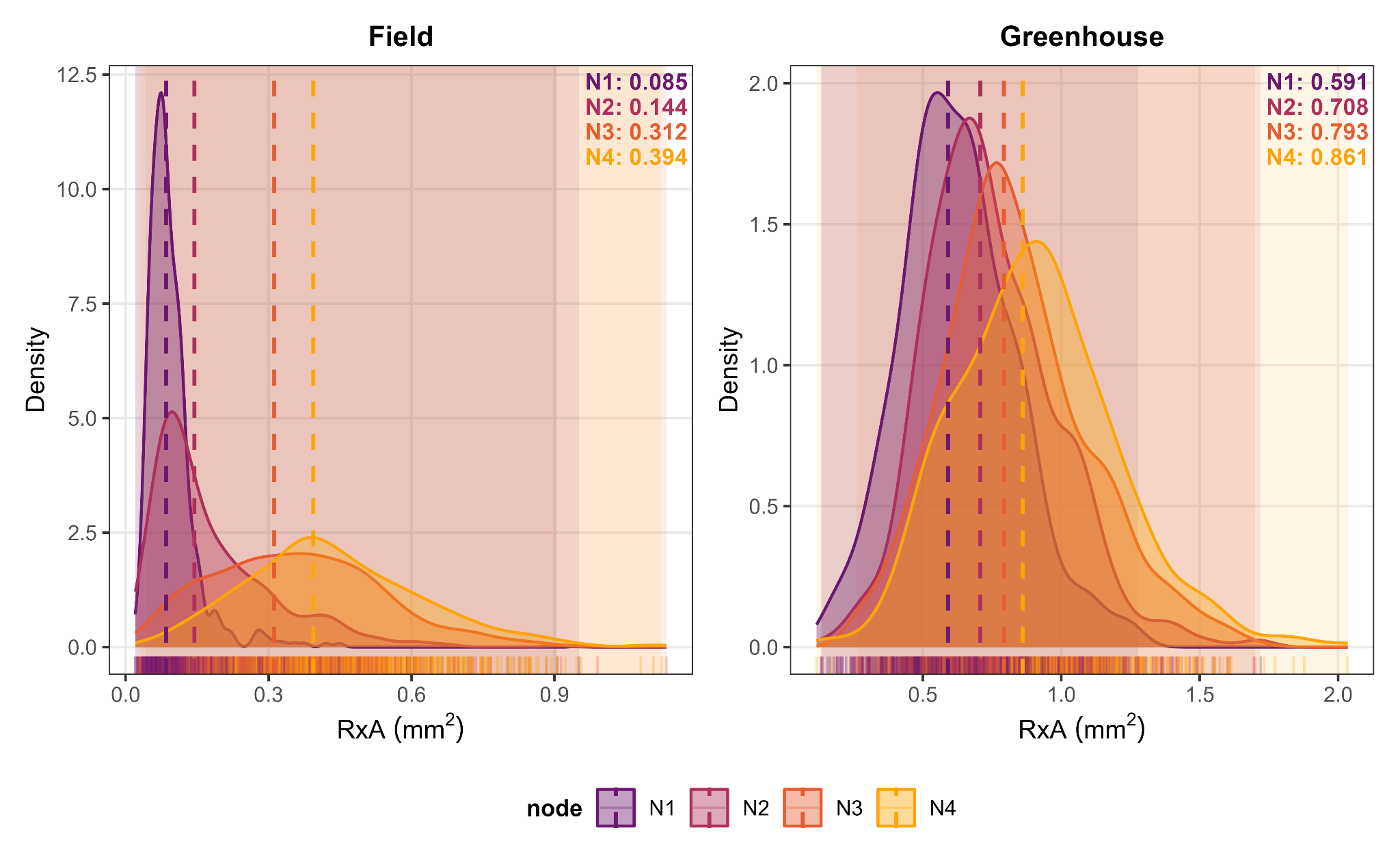

###### **Figure S6. Root cross-sectional area (RxA) density distributions defining the node-specific allometric reference size (X_0,node_) at respective nodal positions.** Density plots with different colours representing nodal positions illustrate the shift in whole-root thickness across sequential nodes in both the field (left panel) and greenhouse (right panel) environments. The faint shaded regions indicate the empirical minimum and maximum range of the RxA for the node. The colored dashed line marks X_0,node_, representing the node-specific environmental baseline RxA, which is used in the allometric size-correction method for genetic analyses.

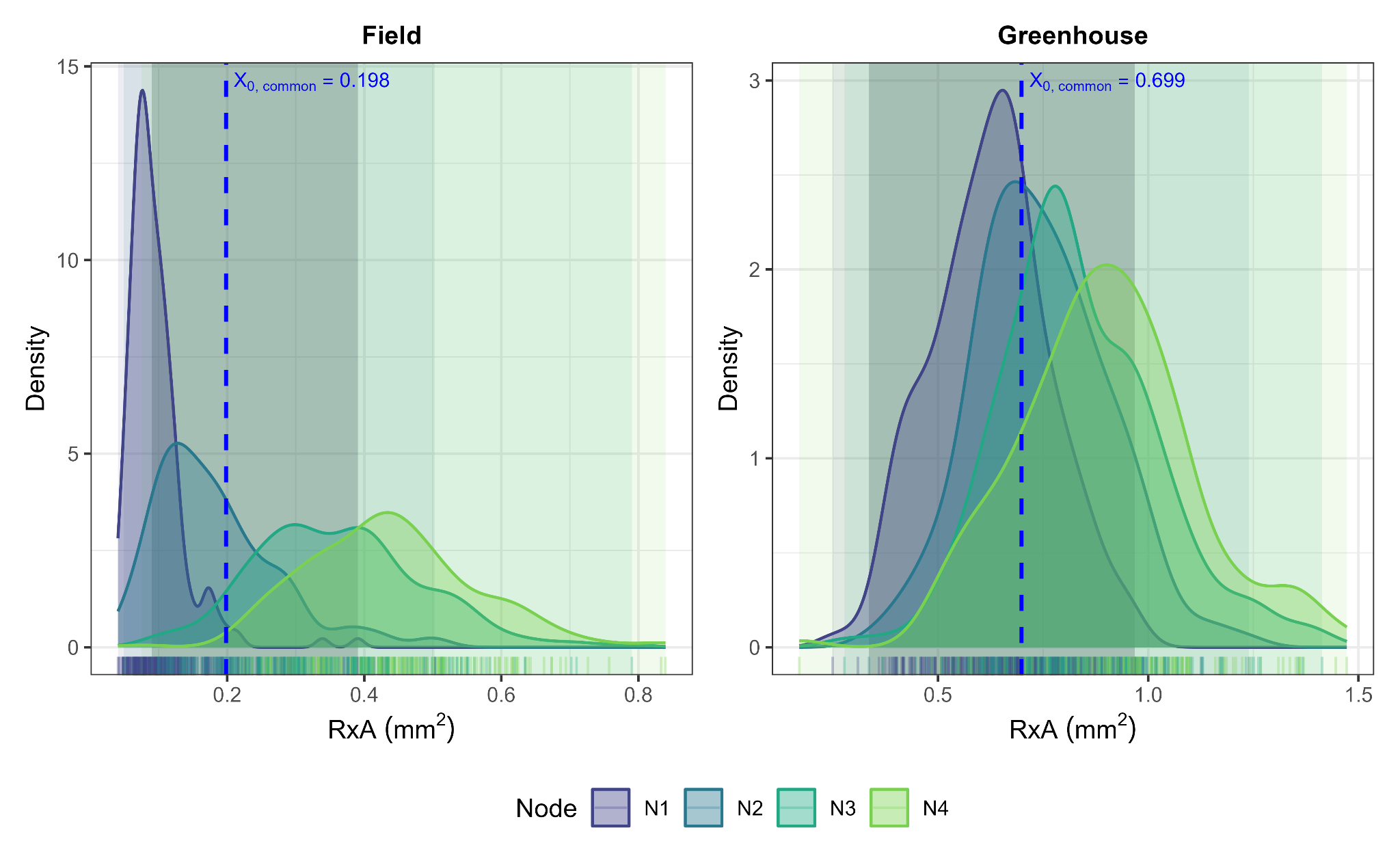

###### **Figure S7. Root cross-sectional area (RxA) density distributions defining the shared reference size (X_0,common_) across N1–N4 nodal positions.** Density plots illustrating the variation in RxA across successive nodal positions in both the field (left panel) and greenhouse (right panel) environments. The dark grey vertical shaded region indicates the empirical overlap in RxA values spanning all four nodes (Field: 0.09 - 0.39 mm^2^; Greenhouse: 0.34 - 0.97 mm^2^). The blue dashed line marks X_0,common_, representing the common environmental baseline root size computed from the overlap region and used in the allometric size-correction method to evaluate physiological gradients.

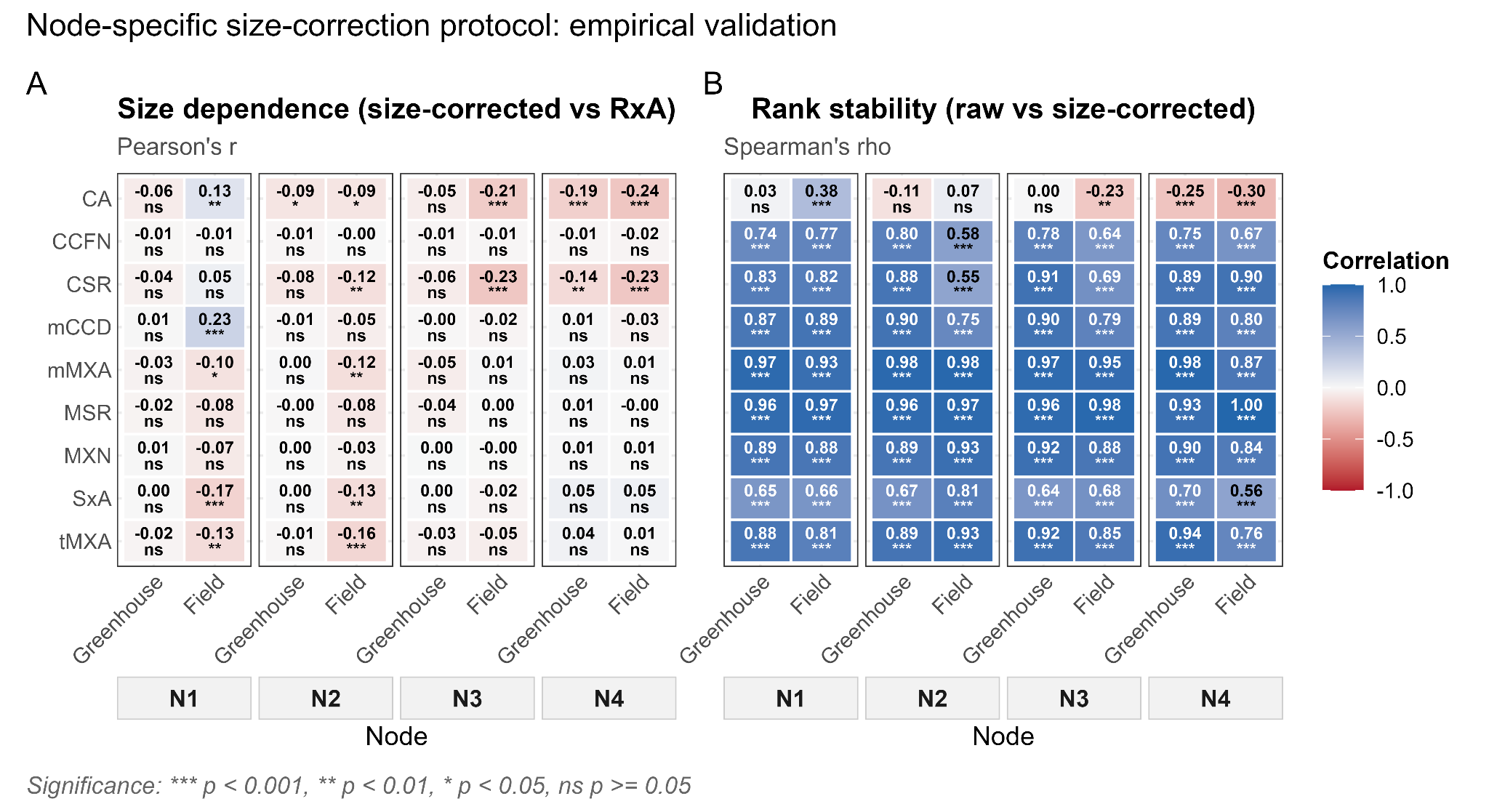

###### **Figure S8. Diagnostic validation of node-specific allometric size correction across diverse environments.** (A) RxA dependence diagnostic using Pearson’s correlation (r) between the allometrically size-adjusted continuous anatomical traits and whole-root cross-sectional area (RxA). (B) Genotype rank stability diagnostic using Spearman’s rank correlation (ρ) between raw and size-adjusted trait entry means. Significance levels for correlations: * *P* < 0.05, ** *P* < 0.01, *** *P* < 0.001, ns = not significant. Trait abbreviations: CA, cortex area; CCFN, cortical cell file layer number; mCCD, mean cortical cell diameter; CSR, cortex-to-stele area ratio (CA/SxA); SxA, stele cross-sectional area; MXN, metaxylem vessel number; tMXA, total metaxylem area; mMXA, mean metaxylem area; MSR, metaxylem-to-stele area ratio (tMXA/SxA).

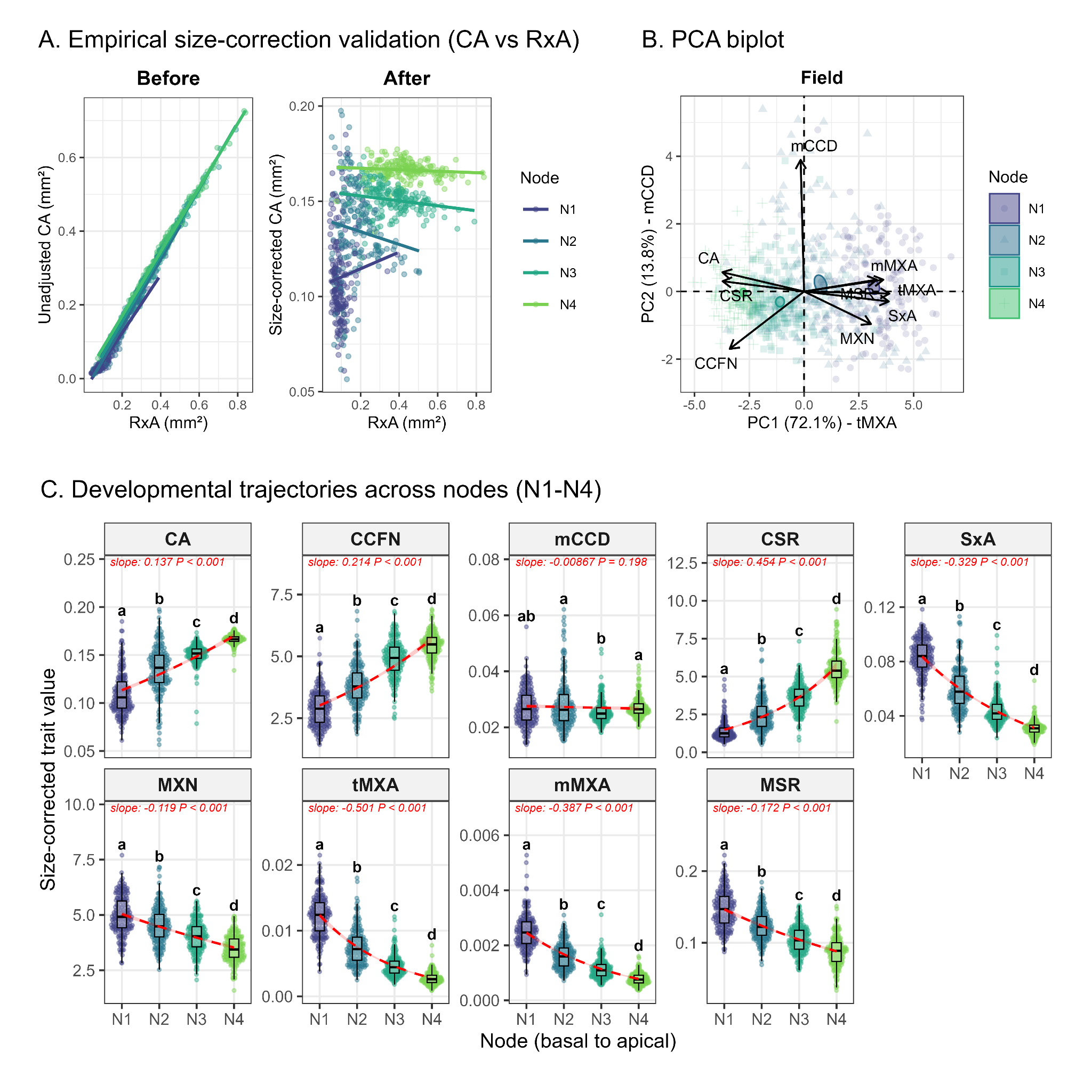

###### **Figure S9. Anatomical variation across successive nodal positions after allometric size correction in field-grown wheat.** (A) Validation of the allometric size-correction approach using cortex area (CA) as a representative trait. Relationships between CA and root cross-sectional area (RxA) are shown before (left) and after (right) allometric size correction. The near-zero slope after correction indicates that the association between CA and RxA was substantially reduced (Figure S11). (B) Principal component analysis (PCA) of size-corrected anatomical traits across four nodal positions. Points are colored by nodal position, ellipses represent 95% confidence regions, and arrows indicate trait loading vectors. Axes show the percentage of variance explained by each principal component. Cortex area (CA), cortical cell file layer number (CCFN), mean cortical cell diameter (mCCD), cortex-to-stele area ratio (CSR = CA/SxA), stele cross-sectional area (SxA), metaxylem vessel number (MXN), total metaxylem area (tMXA), mean metaxylem area (mMXA), and metaxylem-to-stele area ratio (MSR = tMXA/SxA). (C) Variation in size-corrected anatomical traits across four successive nodal positions (N1–N4), standardised to a common reference root cross-sectional area (X_0,common_ = 0.198 mm²; field). All anatomical traits, including continuous traits and count traits (MXN and CCFN), were treated as continuous variables, and post-hoc ratios (CSR and MSR; estimated from their independently size-corrected components) were adjusted using node-specific allometric scaling exponents and analysed using generalised linear mixed-effects models (GLMMs) with a Gamma distribution and log link. Red lines and points indicate model-predicted trends, with slopes and associated P-values shown for each trait. Different lowercase letters indicate significant differences among nodal positions (Tukey's HSD, *P* < 0.05). Trait values are recorded in Table S6.

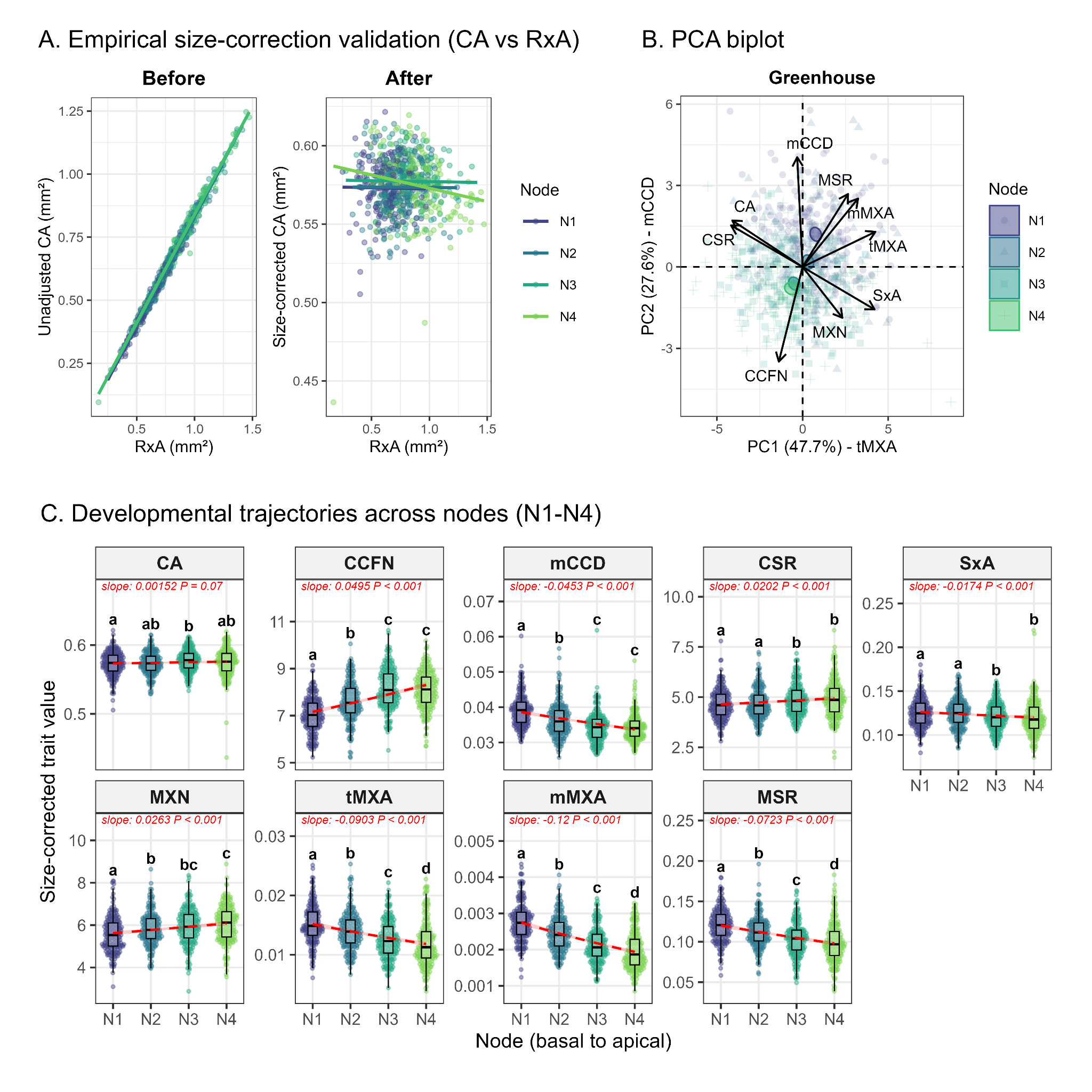

###### **Figure S10. Anatomical variation across successive nodal positions after allometric size correction in greenhouse-grown wheat.** (A) Validation of the allometric size-correction approach using cortex area (CA) as a representative trait. Relationships between CA and root cross-sectional area (RxA) are shown before (left) and after (right) allometric size correction. The near-zero slope after correction indicates that the association between CA and RxA was substantially reduced (Figure S12). (B) Principal component analysis (PCA) of size-corrected anatomical traits across four nodal positions. Points are colored by nodal position, ellipses represent 95% confidence regions, and arrows indicate trait loading vectors. Axes show the percentage of variance explained by each principal component. Cortex area (CA), cortical cell file layer number (CCFN), mean cortical cell diameter (mCCD), cortex-to-stele area ratio (CSR = CA/SxA), stele cross-sectional area (SxA), metaxylem vessel number (MXN), total metaxylem area (tMXA), mean metaxylem area (mMXA), and metaxylem-to-stele area ratio (MSR = tMXA/SxA). (C) Variation in size-corrected anatomical traits across four successive nodal positions (N1–N4), standardised to a common reference root cross-sectional area (X_0,common_ = 0.699 mm²; greenhouse). All anatomical traits, including continuous traits and count traits (MXN and CCFN), were treated as continuous variables, and post-hoc ratios (CSR and MSR; estimated from their independently size-corrected components) were adjusted using node-specific allometric scaling exponents and analysed using generalised linear mixed-effects models (GLMMs) with a Gamma distribution and log link. Red lines and points indicate model-predicted trends, with slopes and associated P-values shown for each trait. Different lowercase letters indicate significant differences among nodal positions (Tukey's HSD, *P* < 0.05). Trait values are recorded in Table S6.

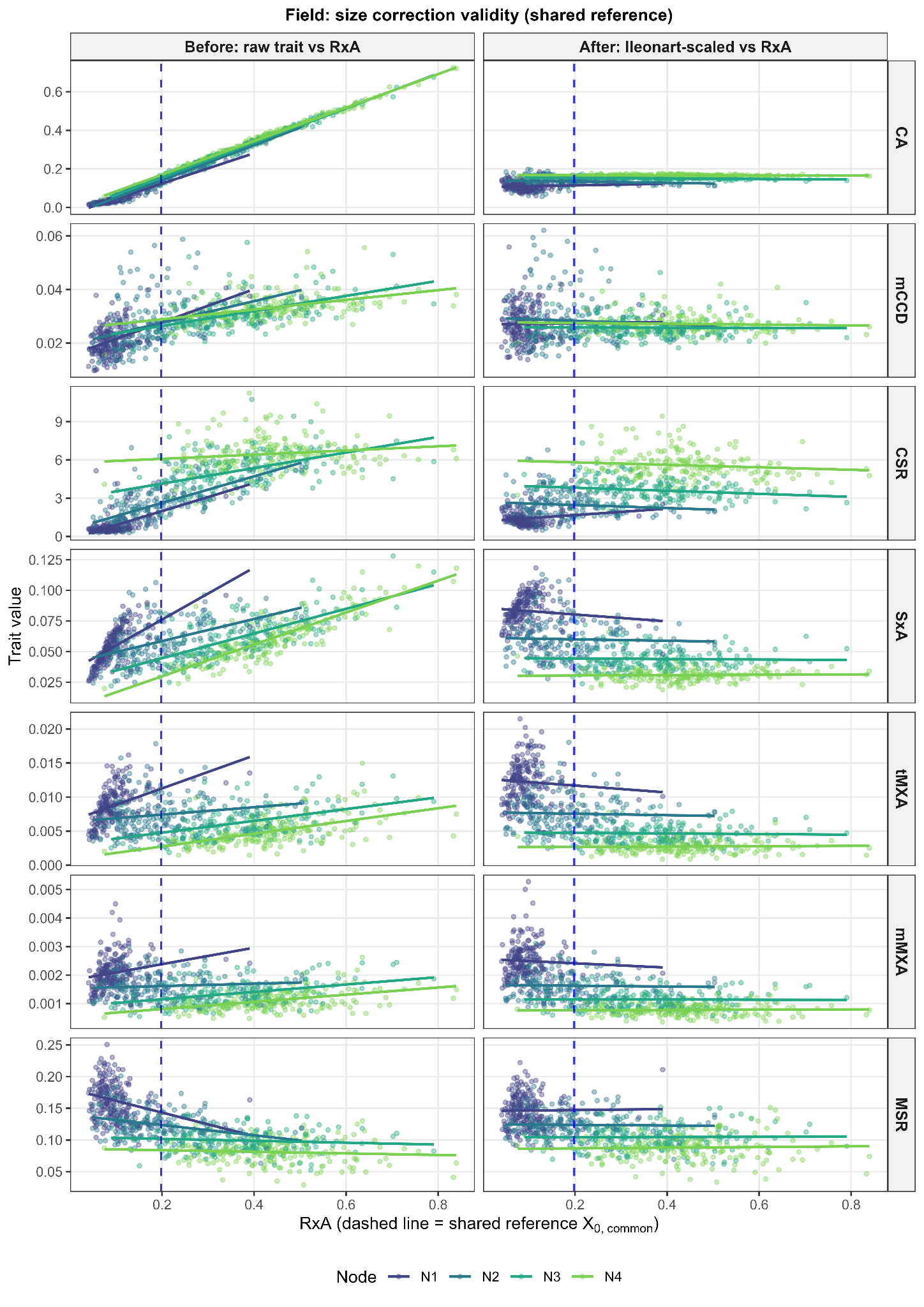

###### **Figure S11. Empirical validation of allometric size correction across anatomical traits and stem nodes N1- N4 for the field dataset.** Left subpanels (Before) show bivariate regressions of raw anatomical trait values against root cross-sectional area (RxA), demonstrating strong positive allometric scaling across nodes N1 through N4. Right subpanels (After) confirm the removal of size dependence post-correction (slope ≈ 0), with values standardised to the shared reference size (X_0,common_, blue dashed line). Separate regression lines for each nodal position were used to assess residual associations between size-corrected anatomical traits and RxA. Trait abbreviations: CA, cortex area; CCFN, cortical cell file layer number; mCCD, mean cortical cell diameter; CSR, cortex-to-stele area ratio (CA/SxA); SxA, stele cross-sectional area; MXN, metaxylem vessel number; tMXA, total metaxylem area; mMXA, mean metaxylem area; MSR, metaxylem-to-stele area ratio (tMXA/SxA).

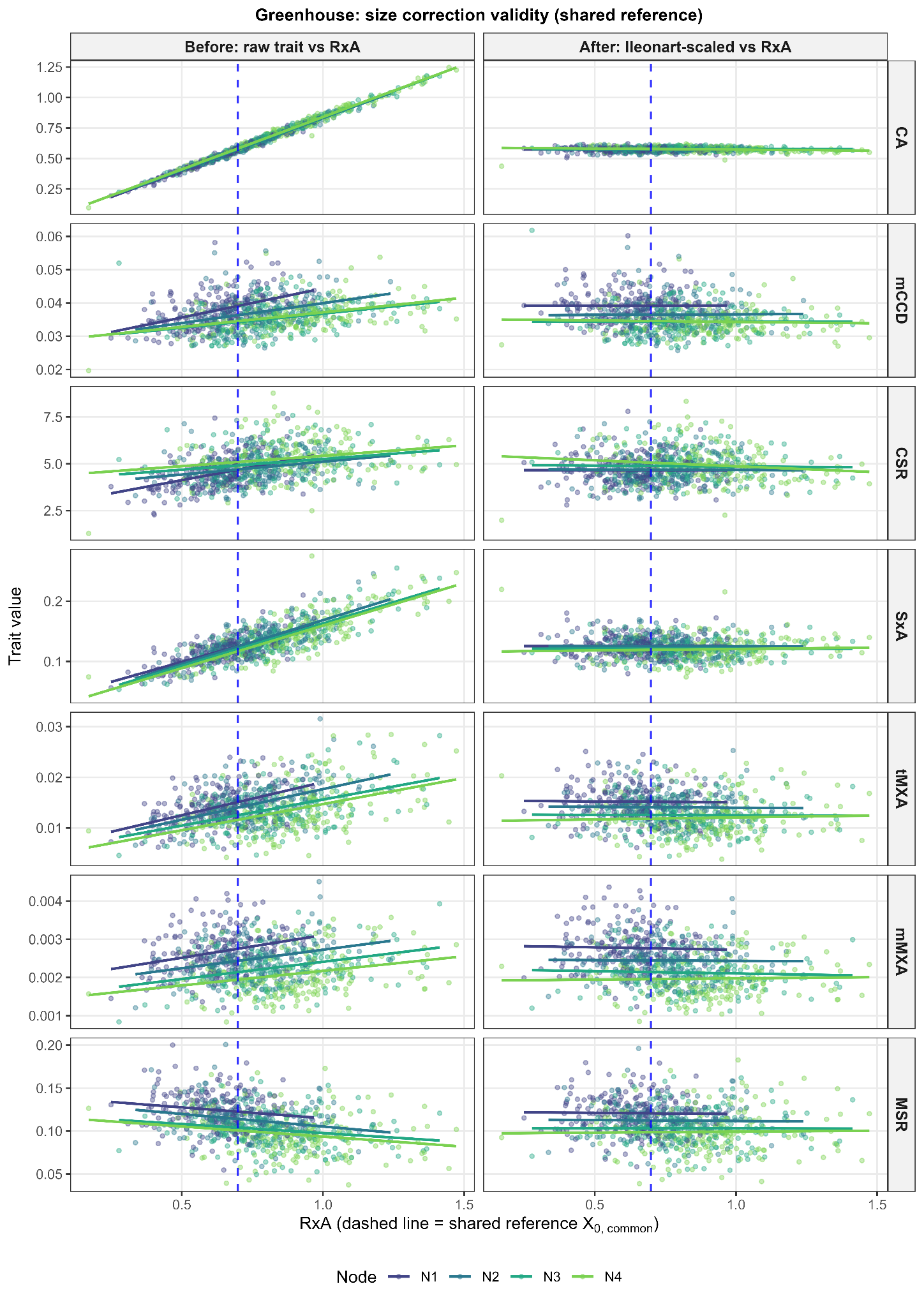

###### **Figure S12. Empirical validation of allometric size correction across anatomical traits and stem nodes N1- N4 for the greenhouse dataset.** Left subpanels (Before) show bivariate regressions of raw anatomical trait values against root cross-sectional area (RxA), demonstrating strong positive allometric scaling across nodes N1 through N4. Right subpanels (After) confirm the removal of size dependence post-correction (slope ≈ 0), with values standardised to the shared reference size (X0,common, blue dashed line). Separate regression lines per node to verify that residual variation is independent of root area. Trait abbreviations: CA, cortex area; CCFN, cortical cell file layer number; mCCD, mean cortical cell diameter; CSR, cortex-to-stele area ratio (CA/SxA); SxA, stele cross-sectional area; MXN, metaxylem vessel number; tMXA, total metaxylem area; mMXA, mean metaxylem area; MSR, metaxylem-to-stele area ratio (tMXA/SxA).

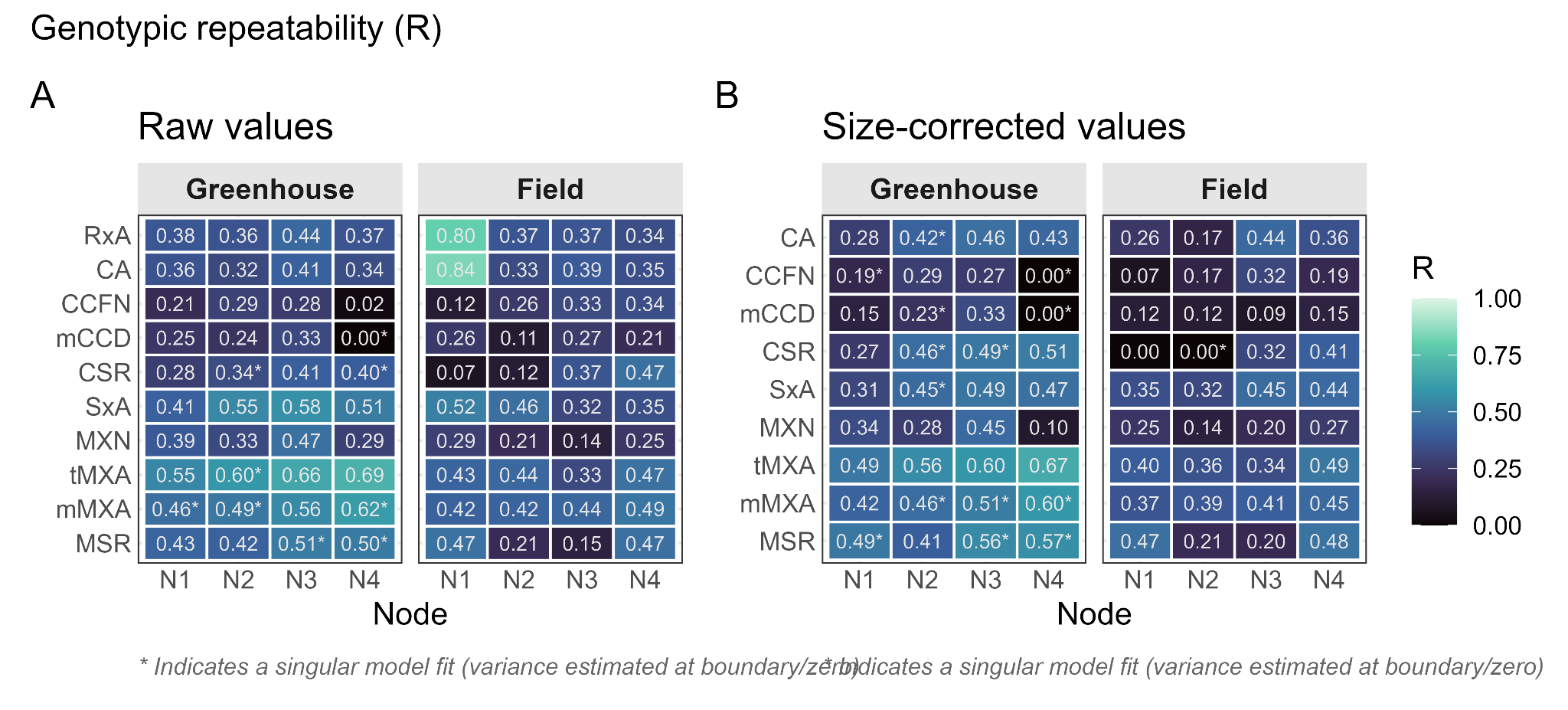

###### **Figure S13. Environment-specific genotypic repeatability (R).** Values estimated for (A) raw and (B) size-corrected trait values via linear mixed models, fit separately for each trait, node, and environment; asterisks indicate singular model fits. RxA appears only in the raw-value panel, as it is the size covariate used for allometric correction. Trait abbreviations: CA, cortex area; CCFN, cortical cell file layer number; mCCD, mean cortical cell diameter; CSR, cortex-to-stele area ratio (CA/SxA); SxA, stele cross-sectional area; MXN, metaxylem vessel number; tMXA, total metaxylem area; mMXA, mean metaxylem area; MSR, metaxylem-to-stele area ratio (tMXA/SxA).

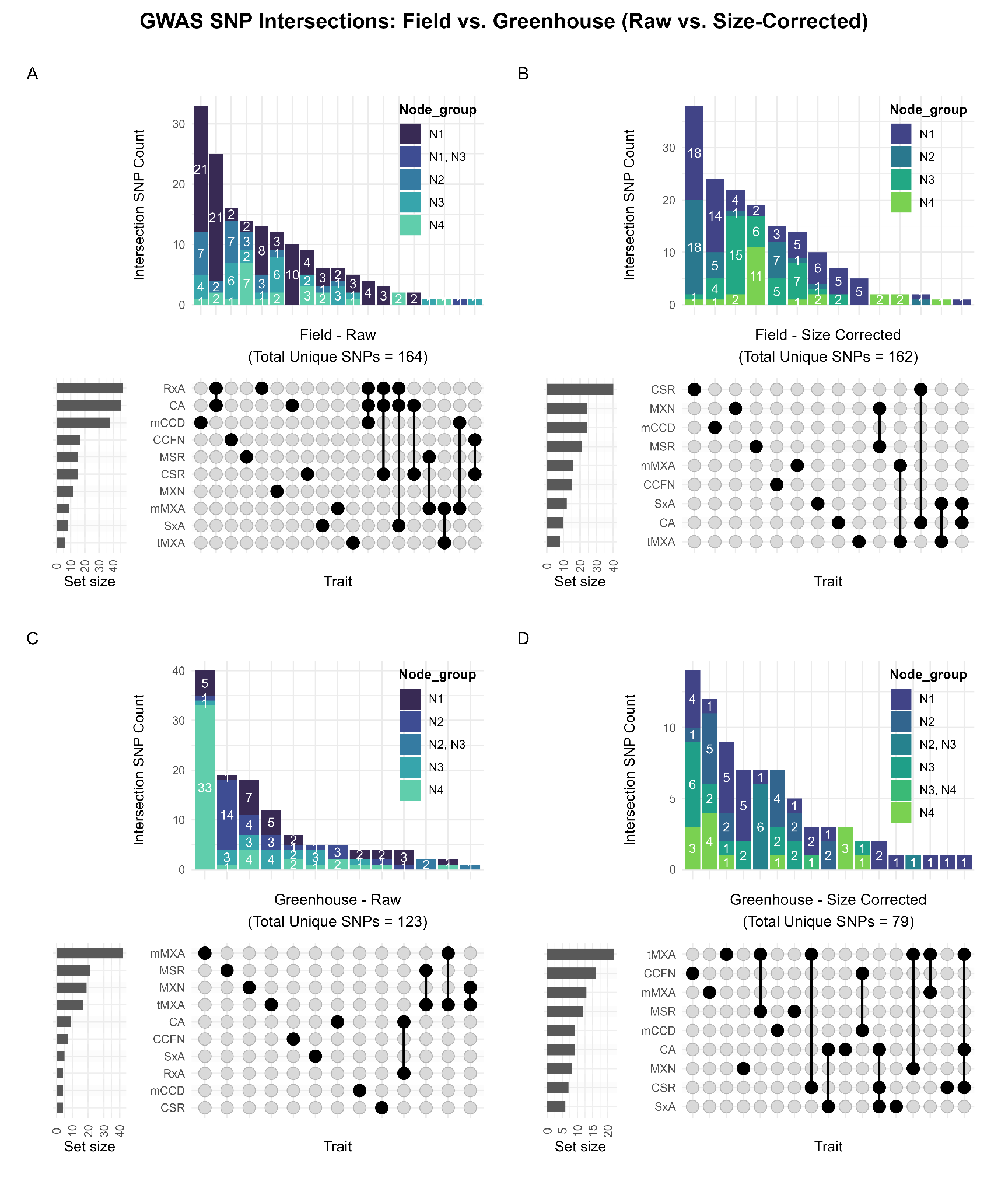

###### **Figure S14. Comparison of genetic associations for unadjusted and size-corrected root anatomical traits across field and greenhouse environments.** UpSet plots illustrate the intersection of suggestive (−log10 *P* ≥ 3.64; FarmCPU model) single-nucleotide polymorphisms (SNPs) across raw, unadjusted anatomical traits (A, C) and allometric size-corrected traits (B, D) evaluated within the field (A, B) and greenhouse (C, D) environments. Vertical bars quantify the number of unique SNPs for a given trait intersection. Bar segments are coloured by the node (or nodal combination) from which the association was mapped, highlighting stage-specific genetic loci. Horizontal bars indicate the total number of suggestive SNPs detected for each individual trait overall. The total number of unique SNPs mapped within each specific environment and methodological pipeline is provided in the respective panel titles. Trait abbreviations: CA, cortex area; CCFN, cortical cell file layer number; mCCD, mean cortical cell diameter; CSR, cortex-to-stele area ratio (CA/SxA); SxA, stele cross-sectional area; MXN, metaxylem vessel number; tMXA, total metaxylem area; mMXA, mean metaxylem area; MSR, metaxylem-to-stele area ratio (tMXA/SxA).

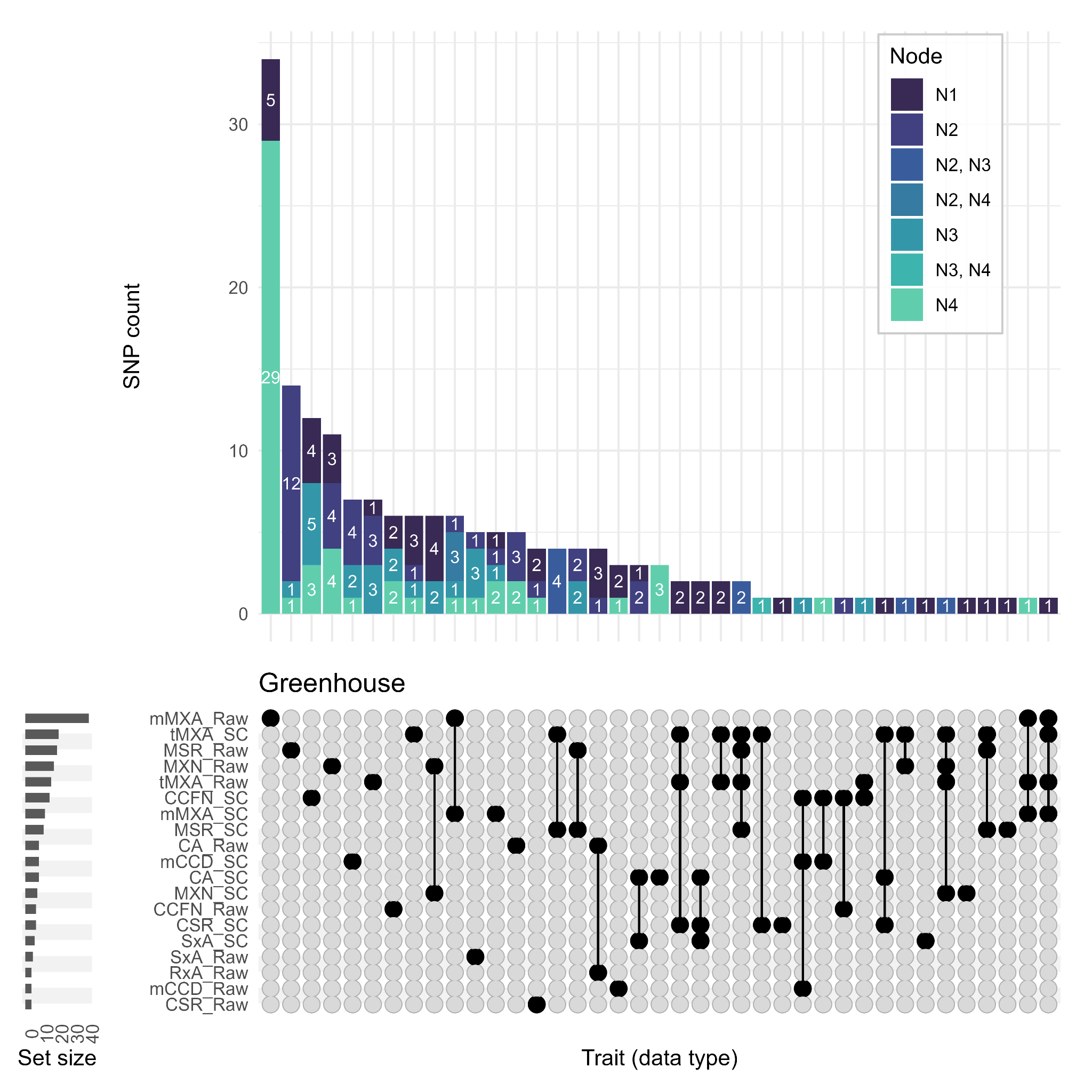

**Figure S15. Shared and unique genetic loci associated with raw and size-corrected root anatomical traits within the greenhouse environment.** UpSet plot illustrating the intersection of suggestive (−log10 *P* ≥ 3.64; FarmCPU model) single-nucleotide polymorphisms (SNPs) across unadjusted (“_Raw”) and allometric size-corrected (“_SC”) traits. Vertical bars denote the number of SNPs unique to a specific trait-version combination or shared across multiple sets. Bar segments are colored according to the specific nodal positions (N1–N4) from which the significant associations were mapped. Horizontal bars display the total number of suggestive SNPs detected for each individual trait version. Trait abbreviations: CA, cortex area; CCFN, cortical cell file layer number; mCCD, mean cortical cell diameter; CSR, cortex-to-stele area ratio (CA/SxA); SxA, stele cross-sectional area; MXN, metaxylem vessel number; tMXA, total metaxylem area; mMXA, mean metaxylem area; MSR, metaxylem-to-stele area ratio (tMXA/SxA).

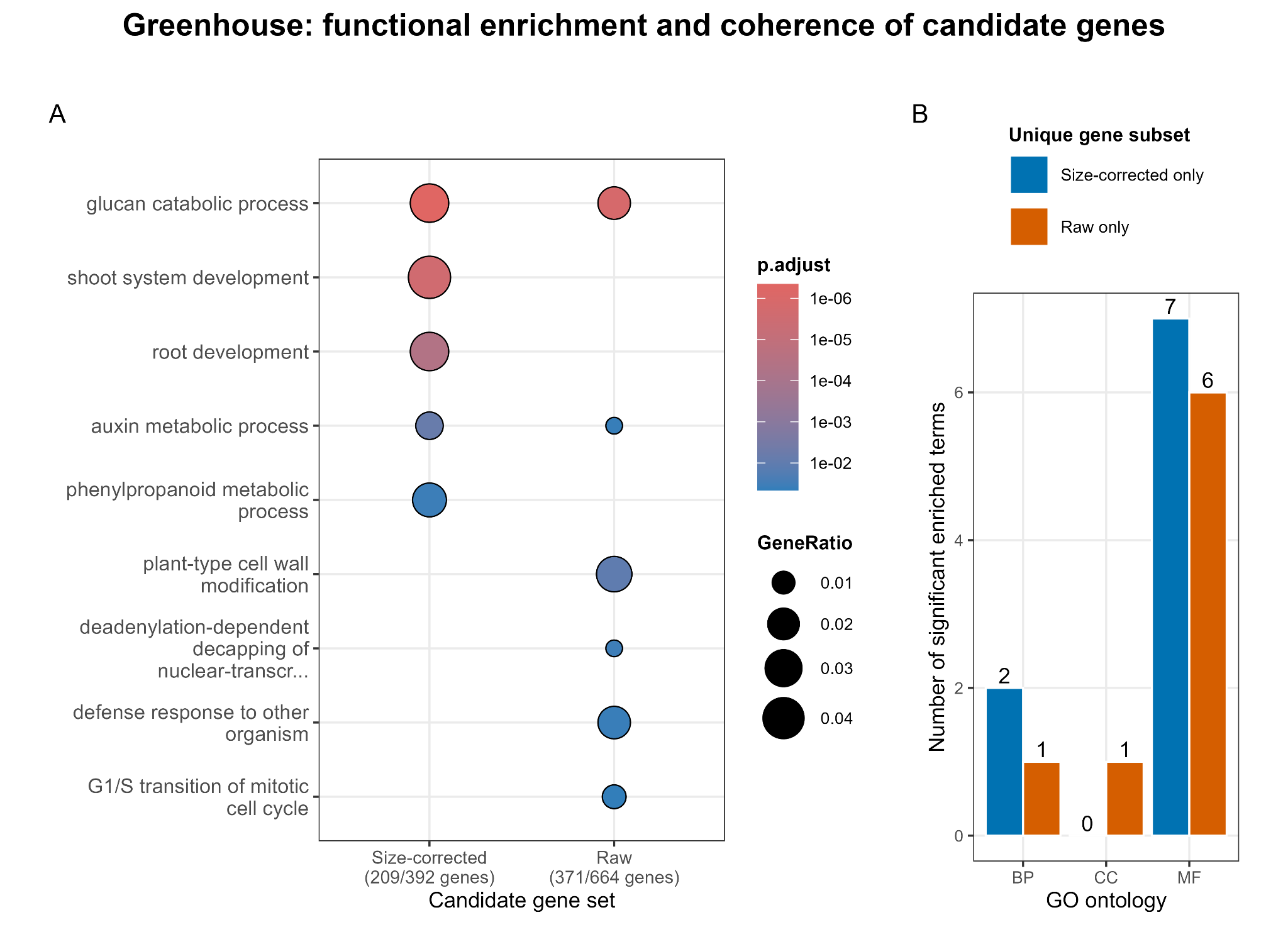

**Figure S16. Functional enrichment and coherence of candidate genes under greenhouse conditions.** (A) Side-by-side dot plot of significantly enriched Biological Process (BP) Gene Ontology terms for the size-corrected (138 genes) and unadjusted raw (219 genes) GWAS models. Node size represents the gene ratio, and colour indicates the Benjamini-Hochberg adjusted p-value. (B) Functional coherence of transformation-specific genes, illustrating the total number of significant GO terms (Biological Process [BP], Cellular Component [CC], and Molecular Function [MF]) derived from candidate genes uniquely identified by either the size-corrected or unadjusted models. The size-corrected unique gene set exhibits higher functional coherence, yielding nearly twice as many significant BP terms as the unadjusted unique set.

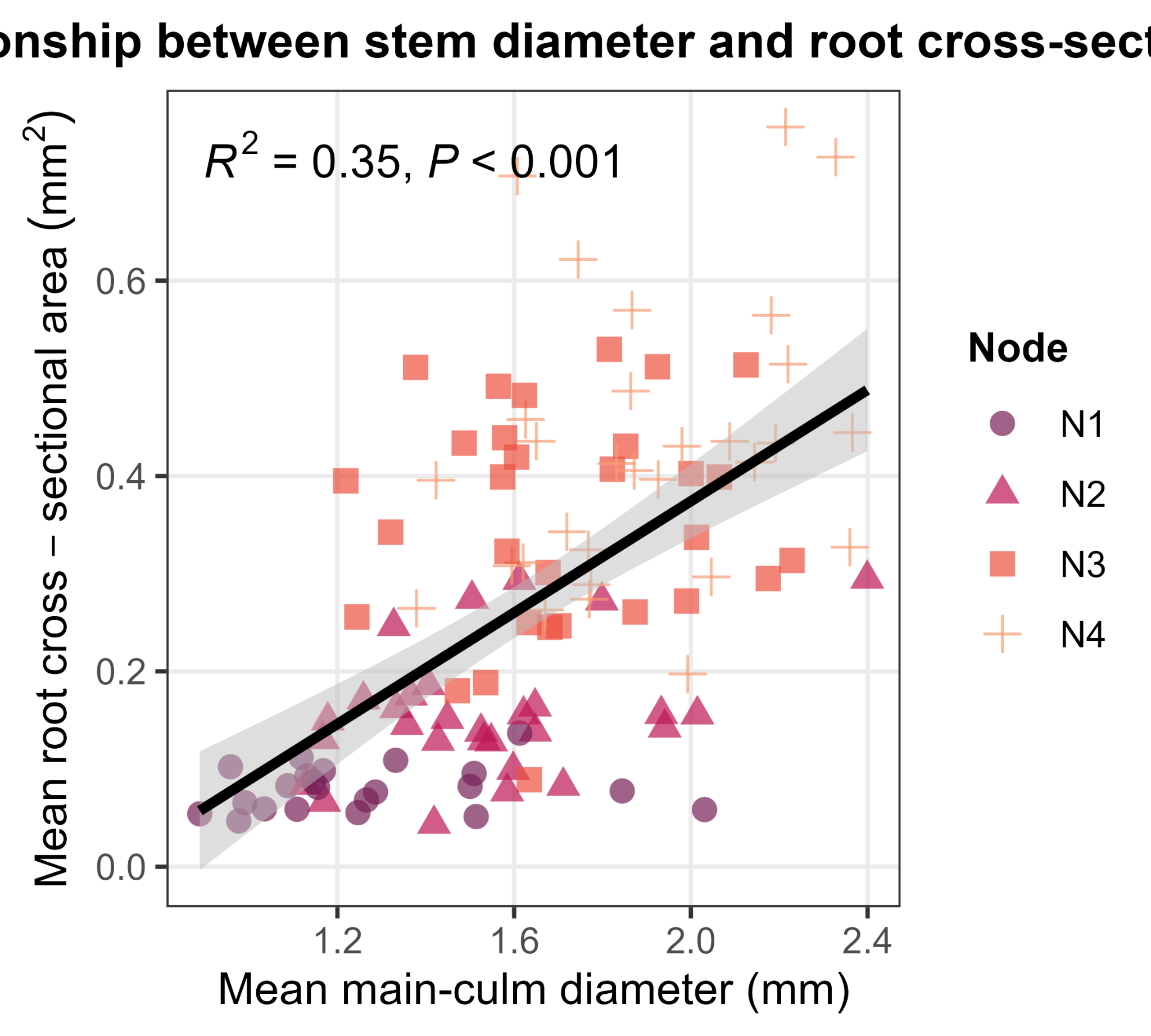

**Figure S17. Scaling relationship between mean main-culm diameter and mean root cross-sectional area across nodal positions.** Ordinary least squares (OLS) regression (R^2^ = 0.35, *P* < 0.001) illustrates the positive association between mean stem diameter and mean root cross-sectional area across 30 wheat genotypes in the field dataset. Data points are colour- and shape-coded by nodal position (N1 through N4, from basal to apical). The solid black line represents the linear regression fit, with the shaded grey band indicating the 95% confidence interval.
